# Accelerating Drug Discovery with HyperLab: An Easy-to-Use AI-Driven Platform

**DOI:** 10.1101/2025.08.31.672525

**Authors:** HyperLab team, Jaechang Lim

**Affiliations:** HITS, 8F, 28, Teheran-ro 4-gil, Gangnam-gu, Seoul, Republic of Korea

**Keywords:** AI drug discovery platform, AI, drug discovery, HyperLab

## Abstract

HyperLab, developed by HITS, is a web-based, AI-driven drug discovery platform designed to increase research efficiency for experimental drug discovery researchers. The platform features an intuitive user interface and experience (UI/UX), enabling researchers without specialized expertise in AI or computational methods to readily generate essential discovery outcomes. By employing a Structure-Based Drug Discovery (SBDD) methodology, HyperLab streamlines the complete discovery workflow. Its core functionalities include the prediction of ligand-protein structures and binding activities (Hyper Binding, Covalent Hyper Binding), protein structure-based molecular optimization (Hyper Design), structure-activity-relationship (SAR) analysis, screening of extensive chemical libraries ranging from one million to seven trillion compounds (Hyper Screening and Hyper Screening X), prediction of ADMET properties (Hyper ADME/T), and an AI-driven assistant designed to boost researcher productivity and efficiency.

In benchmark evaluations, Hyper Binding demonstrated 77% accuracy for binding pose prediction on the PoseBuster v2 benchmark, surpassing conventional docking techniques and closely approaching the 84% accuracy of AlphaFold3, while offering a considerable advantage in computational speed. Furthermore, for binding affinity prediction, HyperBinding showed superior performance over both deep learning and physics-based docking models on two distinct FEP datasets, achieving Pearson correlation coefficients of 0.70 and 0.53, respectively.

For experimental validation, an internal study utilized Hyper Screening to identify top-ranked compounds without without any post-analysis or visual inspection by human experts. The screening process was completed in 24 hours. Through experimental validation of top-ranked compounds, five compounds with IC_50_ values ranging from 70 to 600 nM were identified. Hyper Design was employed to generate novel derivatives with improved predicted binding scores and structural novelty. Of five synthesized compounds, *in vitro* assays confirmed that three demonstrated over 75% inhibition at a 1 µM concentration, with IC_50_ values in the 200 to 400 nM range. Notably, one compound exhibited activity comparable or superior to the reference compound. HyperLab is therefore positioned to substantially lower the barriers of time, cost, and specialized expertise inherent in modern drug discovery initiatives.

## Introduction

HyperLab is an web-based AI-driven drug discovery platform developed by HITS for pharmaceutical researchers. HyperLab is specifically designed for use by all pharmaceuti-cal researchers in drug development, enabling them to leverage its capabilities without needing specialized expertise in computer aided drug discovery. Users can obtain necessary results easily through an intuitive UI/UX, without requiring specialized knowledge or experience in AI or CADD. Key features of HyperLab are predictions of protein-ligand structures and activities, protein structure-based molecular optimization, chemical library screening (from 1 million to 7 trillion compounds), ADME/T prediction, and AI assistant that enhances researcher productivity and efficiency. This technical report explains HyperLab’s main functionalities, underlying technologies, and predictive performance.

## Overview of HyperLab key functionalities

HyperLab has developed state-of-the-art protein structure-based AI models integrated with an intuitive user interface. The primary objective of HyperLab is to facilitate the expeditious identification and optimization of compounds with optimal inhibitory activity against target proteins and variety of pharmacological properties.

HyperLab is an integrated AI platform that streamlines the entire drug discovery pipeline. The workflow can begin with Hyper Binding for accurate protein-ligand binding prediction and affinity assessment. HyperLab also provides optimized worflow for hit identification through Hyper Screening allowing virtual screening of millions of compounds and Hyper Screening X for multi-parameter optimization across trillions of molecules in ultra-large virtual chemical spaces. The platform then transitions to lead optimization using Hyper ADME/T for comprehensive ADMET property prediction and Hyper Design for structure-based molecular optimization to enhance binding affinity while maintaining drug-like properties. To further empower the Hit-to-Lead and Lead Optimization stages, HyperLab incorporates a specialized Structure-Activity Relationship (SAR) analysis module. This integrated approach, supported by an AI assistant throughout the process, significantly reduces the time and resources required for early-stage drug discovery by providing seamless transitions between traditionally fragmented workflows within a single, cohesive environment.

For targets with experimentally determined three-dimensional structures, researchers can initialize the protein structure by specifying a PDB identifier. The proprietary database on the platform enables automatic identification of binding sites and configuration of appropriate binding boxes. Researchers retain the flexibility to manually adjust the locations and dimensions of the binding site as needed. Additionally, the platform supports direct upload of custom PDB files for protein structure initialization. In cases where experimental structures are unavailable, the system can utilize AlphaFold-predicted structures(1) by referencing UniProt identifiers.

Furthermore, HyperLab has implemented advanced Cofolding technology that enables prediction of protein-ligand complex binding conformations directly from protein sequences. This Co-folding methodology significantly streamlines the protein registration workflow by eliminating the need to select specific PDB structures. Researchers need only input the protein name to retrieve sequence data, binding site information, and post-translational modification details from the comprehensive pre-constructed database. Following protein registration, users can modify structural parameters and incorporate multiple proteins to predict binding interactions between protein complexes and ligands. The platform also supports direct input of custom protein sequences independently of the database.

### Hyper Binding

Hyper Binding represents an advanced key function of HyperLab for the prediction of protein-ligand interactions and binding conformations. Hyper binding enhanced generalization performance through the use of a physics-informed deep learning model(2). The system accepts varierty of protein structure input format, including PDB identifier registration, user-uploaded structures, and AlphaFold-predicted conformations as explained in the previous parts. In Hyper Binding, the algorithm explores potential ligand-binding poses within user-defined binding sites while maintaining the protein structure as a rigid entity. Additionally, for cases where only protein sequence information is available, Hyper Binding implements a co-folding approach(3) that simultaneously predicts both the protein structure and ligand binding conformation.

The platform provides comprehensive visualization capabilities, enabling two-dimensional and three-dimensional analysis of predicted binding modes. These structural insights facilitate the mechanistic understanding of protein-ligand interactions and inform rational structure modification strategies to enhance binding affinity. Furthermore, Hyper Binding supports selectivity profiling through comparative binding predictions across multiple protein targets.

Also, Hyper Binding can be utilized to determine the bioassay priority. After uploading a list of candidate molecules designed by medicinal chemists, binding scores can be predicted to establish bioassay priorities. Through cloud-based parallelization, hundreds to thousands of molecules can be rapidly processed, significantly accelerating the computational workflow.

Figure 2. shows the binding mode analysis of Pirtobrutinib (LOXO-305), a selective and non-covalent Bruton’s tyrosine kinase (BTK) inhibitor used in cancer treatment in complex with BTK produced by Hyper Binding. Figure 2-a) displays the three-dimensional representation of Pirtobrutinib bound to the ATP-binding pocket of BTK (PDB ID: 8FLL). The protein is shown as a blue ribbon structure with key in-teracting residues displayed as sticks. The predicted ligand is represented in purple stick format, highlighting its alignment with the X-ray structure represented in cyan stick format. The figure shows the orientation of the ligand within the binding pocket and key interactions with surrounding amino acid residues. Figure 2-b) shows the two-dimensional interaction diagram of Pirtobrutinib with BTK residues calculated using Hyper Binding. The diagram illustrates various interaction types including hydrogen bonds (pink dashed lines), cation-π interactions (yellow dashed lines), and π-π stacking (light pink). Key residues involved in binding include MET477, LEU528, CYS481, and LYS430. The trifluoromethyl pyrazole group, central phenyl ring, and methoxy-substituted benzamide moiety of Pirtobrutinib form multiple favorable interactions within the binding pocket, explaining its potent and selective inhibition of BTK.

**Fig. 1.**
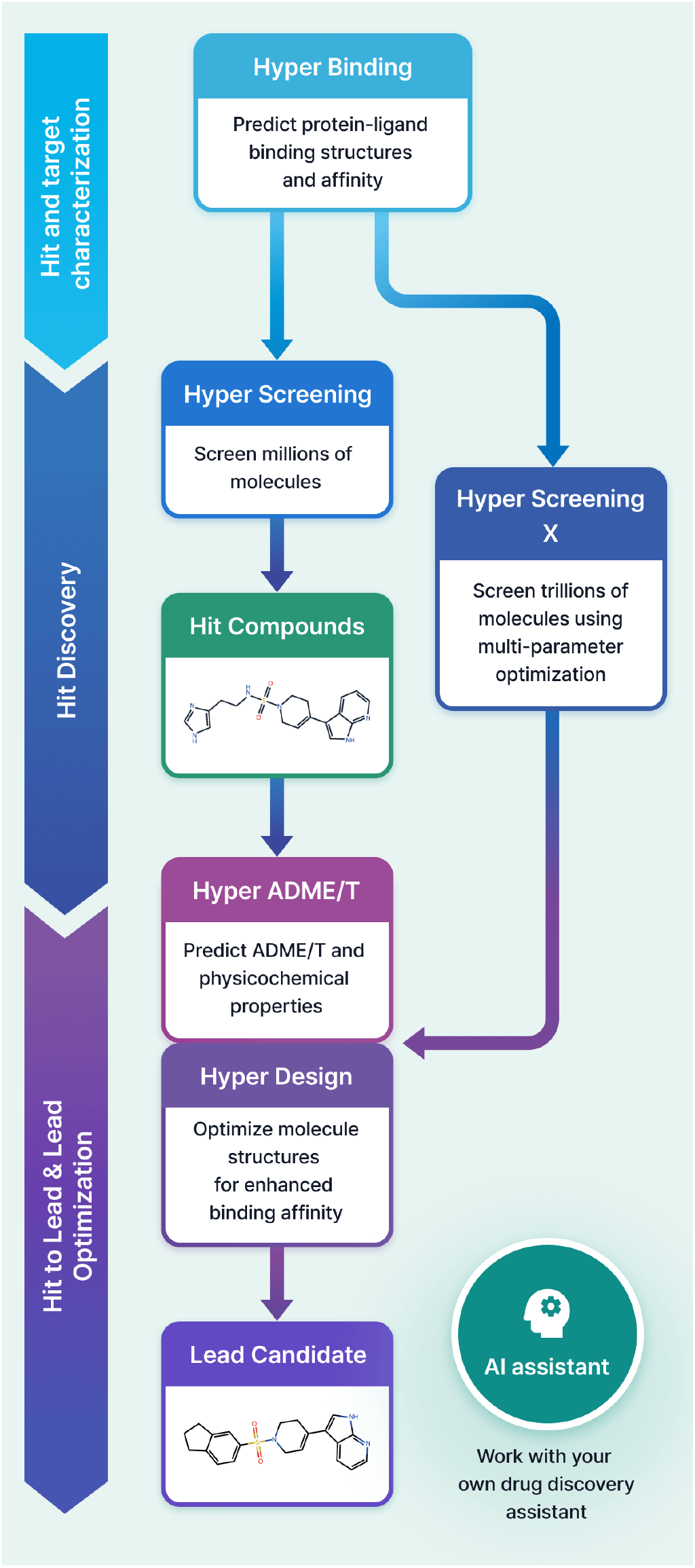
The integrated pipeline demonstrates the comprehensive drug discovery process from hit identification to lead optimization. The workflow progresses through three main phases: (1) hit and target Characterization using Hyper Binding for protein-ligand binding prediction, (2) hit Discovery through virtual screening of millions of compounds and advanced Hyper Screening X for multi-parameter optimization across trillions of molecules, and (3) hit-to-lead and lead optimization employing Hyper ADME/T for ADMET property prediction and Hyper Design for structure-based molecular optimization. The AI assistant provides continuous support throughout the entire discovery process.

**Fig. 2.**
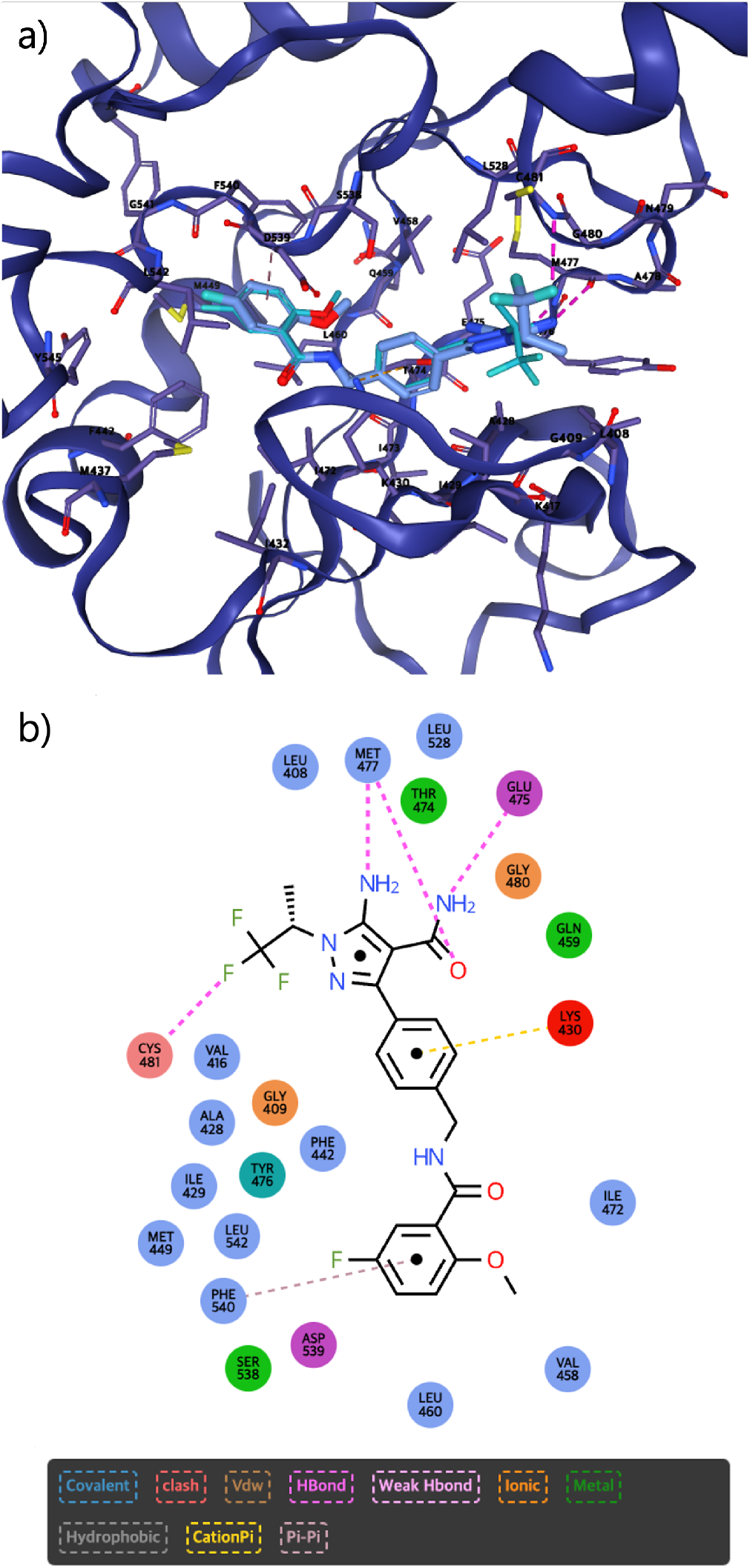
Visualization of Pirtobrutinib (LOXO-305) in complex with Bruton’s tyrosine kinase: (a three-dimensional structural representation and (b two-dimensional interaction diagram.

**Fig. 3.**
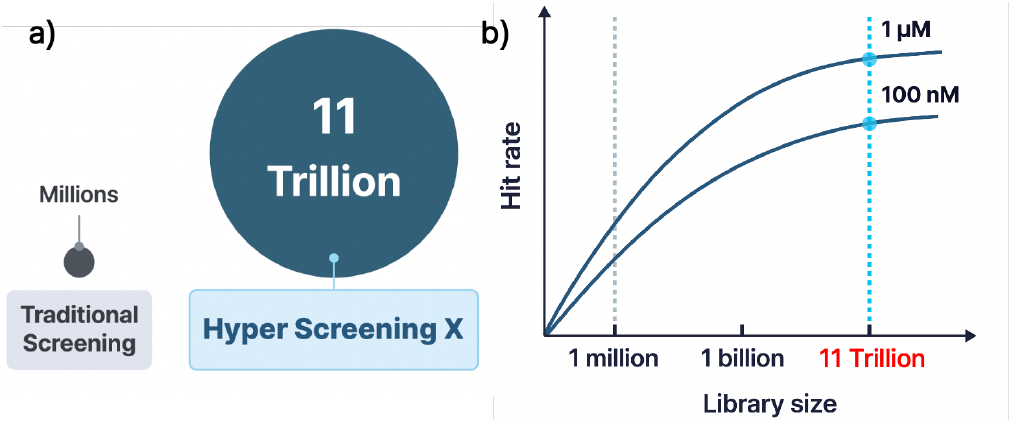
a) Dramatic difference in library size between a conventional docking (millions of compounds) and Hyper Screening X (11 trillion compounds). b) Demonstrates the relationship between library size and hit rate. The graph shows the schematic representation that the increase of the library size to 7 trillion compounds significantly enhances the probability of identifying active compounds.

### Hyper Screening

Hyper Screening is a virtual screening function designed to discover initial active compounds in the early-stage of drug discovery. It performs Hyper Binding calculations on all molecules in a library against a specified protein, then provides the top 500 molecules with the best binding scores. Five pre-processed libraries are available for specific purposes. When Hyper Screening is performed using the provided libraries, compound information and purchase links are provided to enable users to acquire these substances. Users can also register their own libraries to perform screening. Once a library is registered, it can be used for screening against different targets. For the selected top 500 molecules, users can perform 2D/3D interaction analysis and optimize molecular structures using Hyper Design, just as with regular Hyper Binding calculations.

The Table **??** summarizes the five primary molecule libraries available in Hyper Screening. The Fragment library contains 500,000 molecules selected from the Diverse library that meet specific physicochemical criteria (MW < 300, LogP < 3.0, < 4 hydrogen bond donors). The comprehensive Diverse library includes 1 million diverse compounds filtered according to HITS’s proprietary criteria, including Lipinski rule compliance and clustering based on in-stock availability. The FDA-approved library comprises 1,100 marketed drugs, while the Natural product-like fragments library contains 4,200 molecules representing 1,000 structural motifs derived from natural products. The Kinase-focused library of 65,000 compounds includes bioisosteric derivatives of allosteric kinase inhibitors and hinge binder molecules specifically designed for kinase-targeted drug discovery project.

### Hyper Screening X

Hyper Screening X is a service designed to discover active compounds by screening an extensive virtual library of 11 trillion molecules. Compared to conventional virtual screening methods that typically explore around 1 million compounds, Hyper Screening X offers two significant advantages: 1) increased hit discovery probability that can be dozens times higher than traditional approaches(4), and 2) an increased likelihood of identifying novel active compounds compared to conventional commercial screening libraries. Previous research has demon-strated that virtual libraries containing billions of molecules can achieve hit discovery success rates of 20-30%(4). By leveraging a vast chemical space of 11 trillion molecules, Hyper Screening X provides a significantly higher probability of discovering novel, potent hit compounds compared to conventional screening methods that are limited to conventional screening libraries.

**Table 1.**
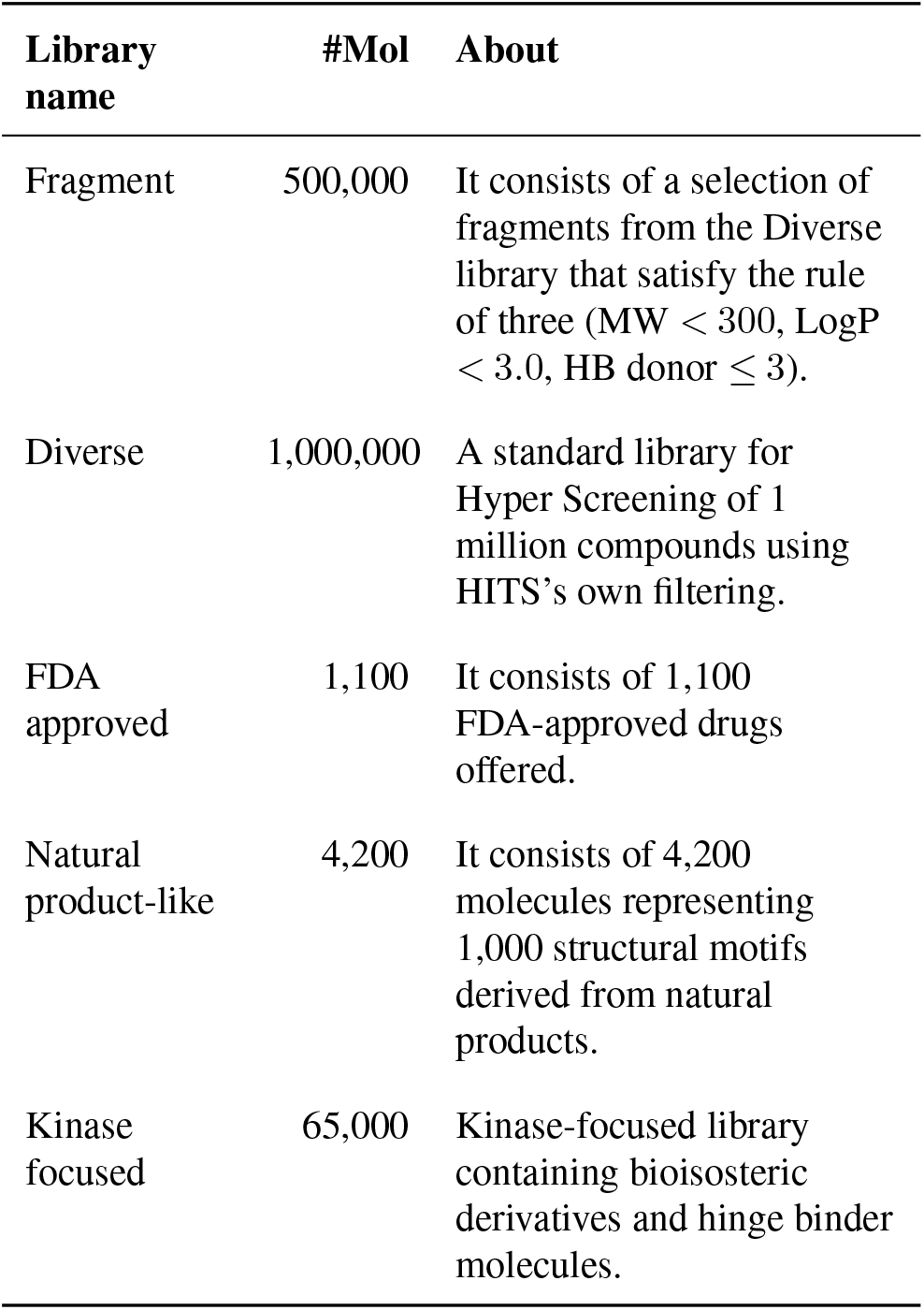
Overview of Hyper Screening libraries.

| Library name | #Mol | About |
| --- | --- | --- |
| Fragment | 500,000 | It consists of a selection of fragments from the Diverse library that satisfy the rule of three (MW < 300, LogP < 3.0, HB donor ≤ 3). |
| Diverse | 1,000,000 | A standard library for Hyper Screening of 1 million compounds using HITS's own filtering. |
| FDA approved | 1,100 | It consists of 1,100 FDA-approved drugs offered. |
| Natural product-like | 4,200 | It consists of 4,200 molecules representing 1,000 structural motifs derived from natural products. |
| Kinase focused | 65,000 | Kinase-focused library containing bioisosteric derivatives and hinge binder molecules. |

Calculating binding affinities for all 11 trillion molecules individually would require astronomical computational resources. To address this challenge, Hyper Screening X employs generative AI to identify combinations of building blocks and reaction templates capable of producing molecules with desired binding scores and properties by the user. Beyond binding affinity, the system can optimize various molecular properties including molecular weight (MW), topological polar surface area (TPSA), and LogP. Hyper Screening X utilizes state-of-the-art GFlowNet(5)-based deep learning models(6) for virtual library exploration, offering the advantage of discovering more diverse molecular structures compared to reinforcement learning-based methods. Additionally, the deep learning models employed by Hyper Screening X demonstrate property optimization performance that is 5-30% superior to similar competitive models(6).

The Hyper Screening X workflow consists of three main stages: 1) target protein and target property setting, 2) generative AI training, and 3) molecule generation upon completion of training. Training the generative AI models takes approximately 48 hours, and the subsequent generation of 100 molecules takes 30 minutes. Following molecule generation, users can request synthesis of the generated compounds through Synple Chem. After consultation with Synple Chem, the synthesized compounds can be delivered to the user. For researchers who prefer to conduct synthesis independently, Hyper Screening X also provides synthetic routes for the generated molecules. Hyper Screening is suitable when a user wants to quickly discover active compounds from their proprietary compound library or from approximately one million commercially available compounds, and then proceed to the hit-to-lead phase. If a user needs a higher hit discovery rate, simultaneous optimization of multiple properties, and identification of novel active compounds, Hyper Screening X is the appropriate choice.

### Hyper Design

Hyper Design provides a feature that transforms given scaffolds to design molecules predicted to show improved biological activity against the target protein. The Hyper Design starts with predicting binding scores and binding conformations of scaffolds, then leverages this information to design molecular structures with improved binding scores. Also, Hyper Design can initiate the process from X-ray structures of small molecule-bound proteins. By registering a protein structure via its PDB ID and selecting the specific site of the ligand for modification, the system recommends new derivative structures. This approach provides inspiration for designing novel substances based on existing ligands.

Users have flexible design options, including the ability to select specific substructures of the scaffold for modification or add fragments at specific atomic positions. During molecular design, Hyper Design considers synthesizability of the generated molecules while replacing fragments or adding new ones. The fragment set used by Hyper Design was derived from the ChEMBL(7) compound list. In addition, the Hyper Design provides 3D structures of generated molecules, enabling users to interpret results and gain insight for further molecular design ideas.

Molecules generated through Hyper Design can be recursively subjected to further design iterations. The resulting molecules are visualized in a hierarchical flow diagram, making it easy to understand the molecular design progression. Hyper Design can serve multiple purposes, including patent circumvention of reference compounds, fragment growth in fragment-based drug design, and generation of novel molecular design ideas. You can find a case study using Hyper Design in the Case Study section.

Figure 4 illustrates an application of HyperLab’s Hyper Design capability in medicinal chemistry, specifically demonstrating hit optimization starting from a small fragment. The top-left structure shows the reference compound Imatinib, while the remaining structures represent novel derivatives generated through Hyper Design. The blue-highlighted portions in each molecule indicate the common hit fragment that served as the starting point for the design process. This scaffold acted as the initial hit molecule in the actual development process of the imatinib molecule(8). This visualization effectively demonstrates how Hyper Design can successfully optimize an initial hit molecule into more complex molecules with structural features similar to known active compounds like Imatinib. The designs build upon the core fragment while exploring different substitution patterns and functional groups that could enhance target binding and pharmacological properties. Each variant maintains the essential hit fragment while introducing strategic modifications that could potentially improve binding affinity against the target.

**Fig. 4.**
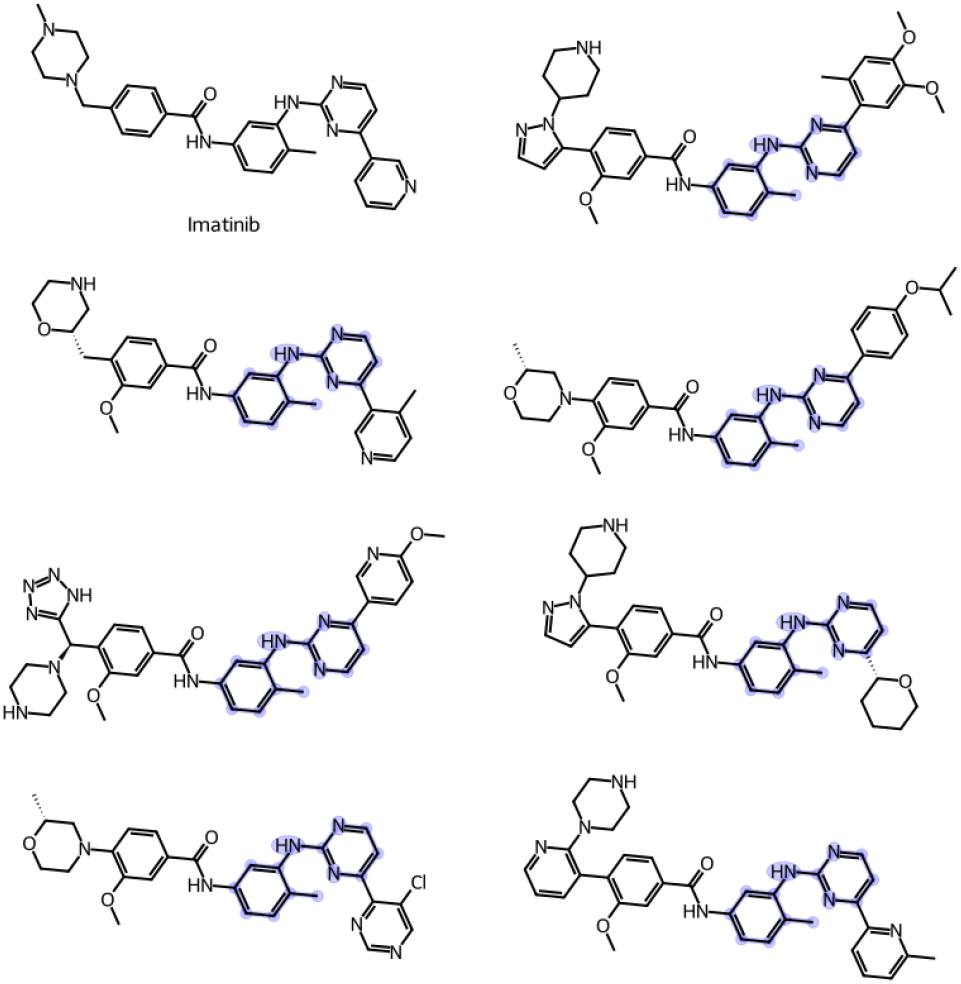
Hit optimization demonstration using Hyper Design: Starting from a hit fragment (highlighted in blue), HyperLab successfully generated multiple Imatinib-like molecules by optimizing the initial hit. This capability enables efficient hit-to-lead development for drug discovery applications.

**Fig. 5.**
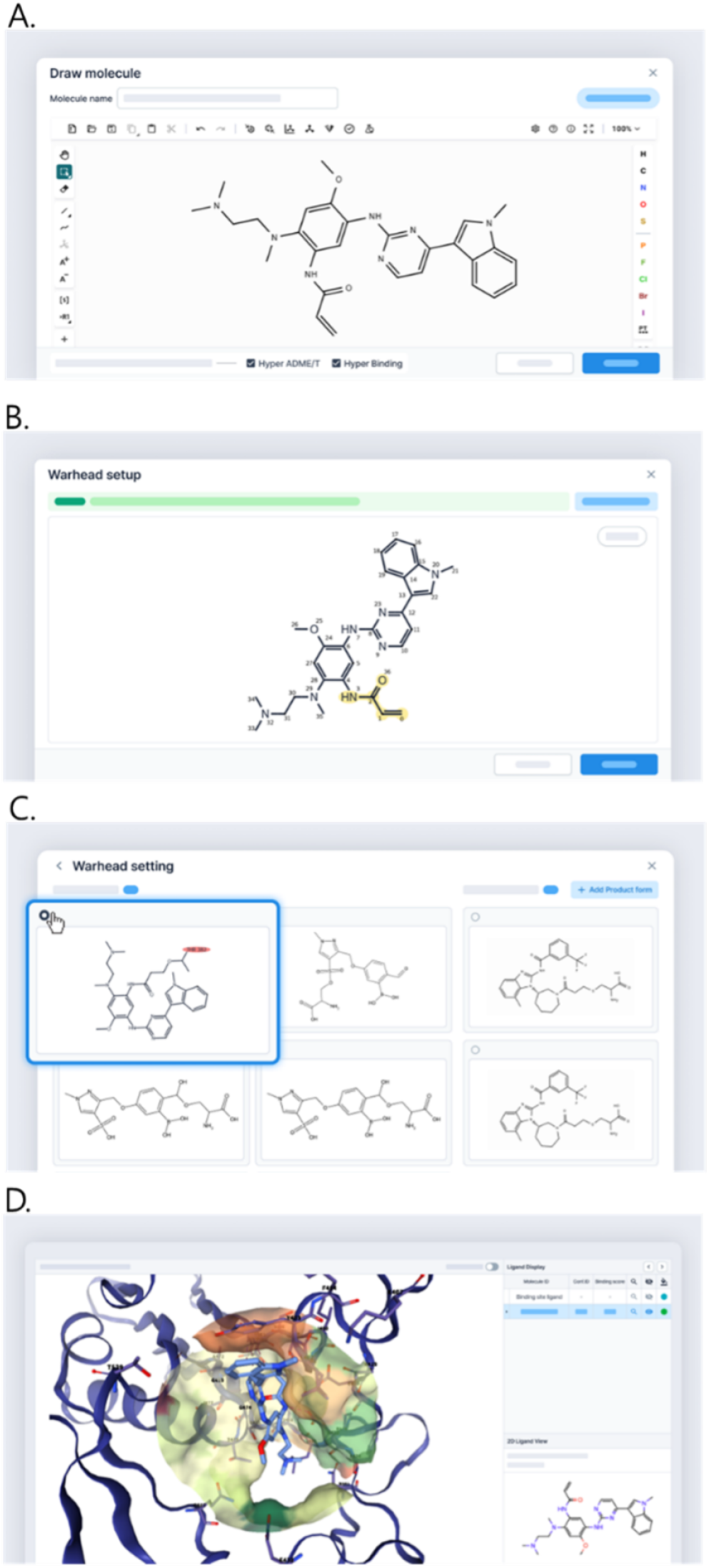
The figure illustrates the Covalent Hyper Binding workflow, which proceeds from molecule registration and warhead selection to the automated recommendation of compatible product forms, and finally to the evaluation of binding scores and 3D predicted poses.

### Covalent Binding, Design and Screening Features

HyperLab provides a comprehensive suite of features for covalent drug discovery, encompassing binding structure prediction, molecule design, and virtual screening. Since the success of covalent binding prediction relies heavily on accurately designating target residues and matching them with viable reactive warheads, HyperLab offers both an Auto-Recommendation mode and a Manual Definition mode. The Auto-Recommendation mode utilizes an internal reaction database to automatically identify compatible warhead-residue pairs and suggest predicted products, significantly streamlining complex configurations that are typically cumbersome in other software. Conversely, the Manual Definition mode allows users to custom-design reactant and product definitions based on specific residues, delivering results in a consistent, warhead-centered format.

These predictions can be performed using either fixed protein structures or by predicting the protein structure directly from the sequence as part of the workflow. Furthermore, this module is tightly integrated with HyperLab Bench’s SAR analysis and Hyper ADMET prediction, providing identical profiles for both reactant and covalent product states so that biological and pharmacological changes can be compared on a single screen. For molecule structure design, Hyper Design allows users to refine the molecular structure by modifying parts of the molecule other than the warhead and leaving group substructure, all while maintaining the covalent bond. Finally, the screening module serves as an extension of covalent docking, utilizing a curated HITS internal database of residue-specific warhead libraries to systematically identify candidates from commercial or private compound collections.

### SAR analysis

HyperLab’s SAR analysis is a specialized module designed to streamline Structure-Activity Relationship (SAR) analysis, which is critical during the Hit-to-Lead and Lead Optimization stages. This feature is seamlessly integrated with “Bench”, allowing users to instantly load molecular lists and begin analysis simply by specifying a Bench when creating an analysis page.

The central component of this analysis is setting a “Core” to group molecules. The system offers “Automatic Recommendation,” which uses BRICS-based fragmentation to suggest core candidates based on common Murcko scaffolds, and “Manual Designation,” where users can directly draw structures. Users can prioritize up to five cores for analysis. Figure 6 illustrates the resulting R-group decomposition table. The moleculess are separated into columns by the designated Core and substituents (R1, R2, etc.) and arranged according to set alignment rules. This consistent view allows users to identify structural differences at a glance and enhances the efficiency of comparative analysis. The analysis is further deepened by the unique “Hierarchical Decomposition” feature. For example, within a complex structure, users can recursively analyze a specific sub-fragment (e.g., designating a secondary core within a substituent) to explore structural diversity in greater depth.

**Fig. 6.**
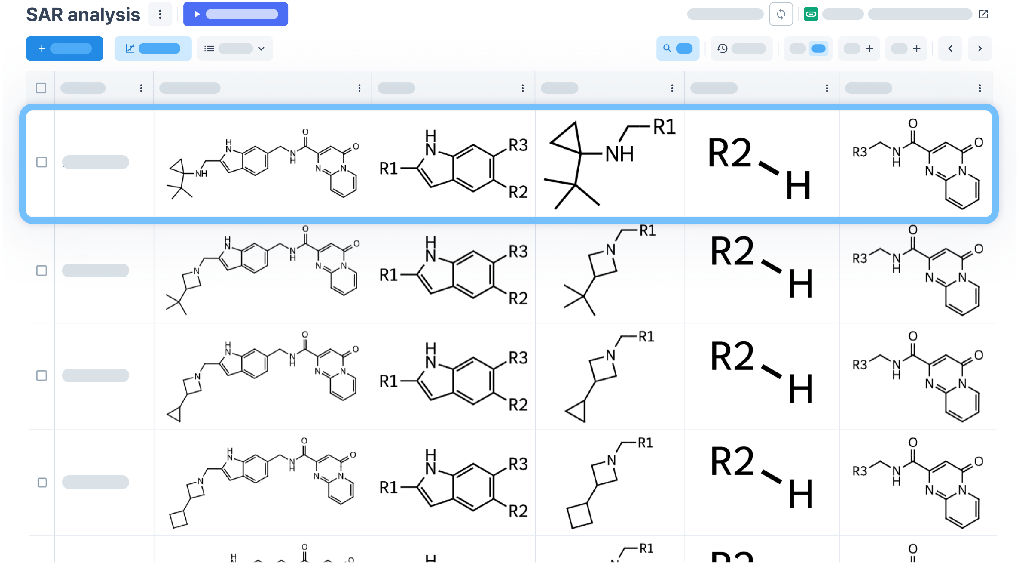
R-group decomposition interface for SAR analysis. The molecules in the library are structurally aligned and decomposed based on the user-designated core. Each column (Core, R1, R2) maintains a consistent 2D alignment, enhancing the efficiency of visual comparative analysis.

**Fig. 7.**
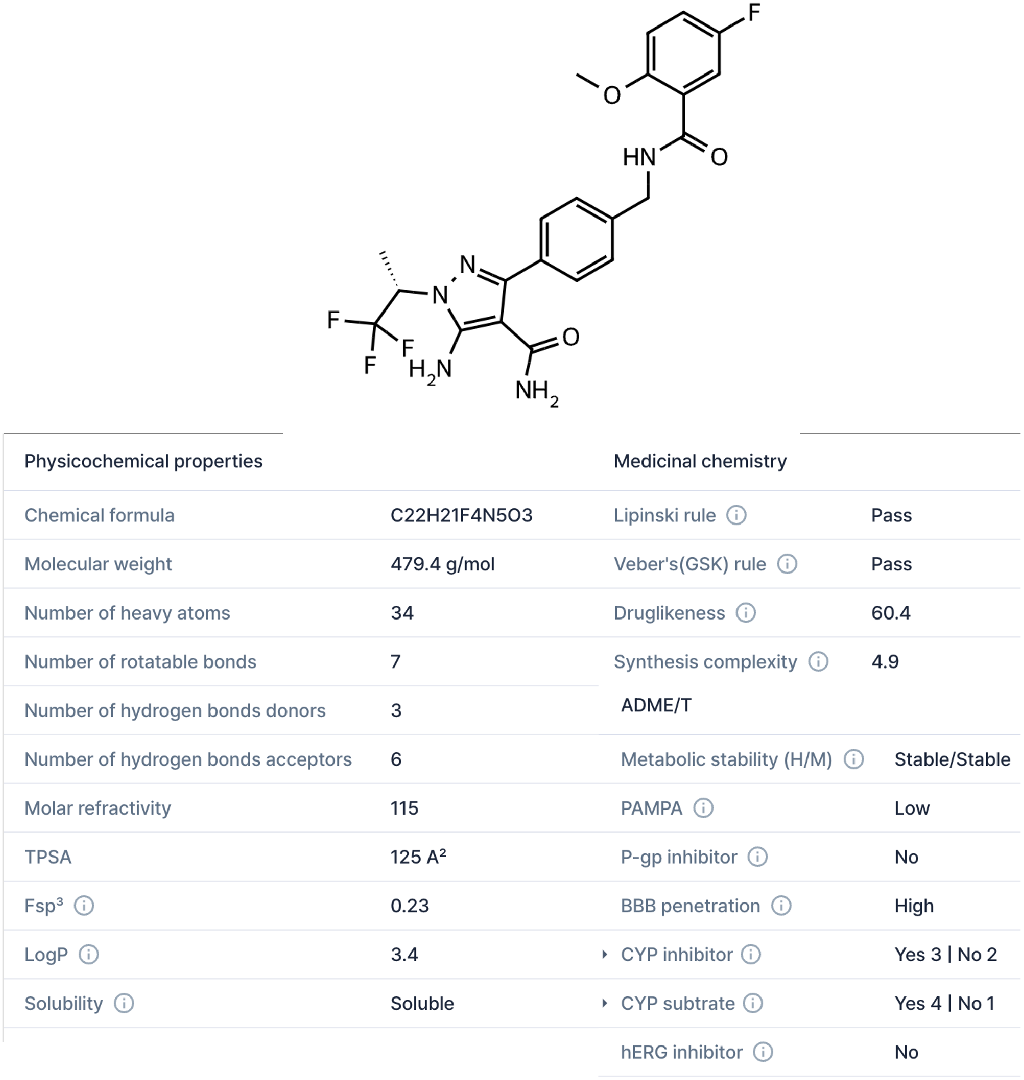
Physicochemical properties, medicinal chemistry properties, and ADME/T profile of Pirtobrutinib predicted by Hyper ADME/T.

In addition, the system integrates HyperLab’s predicted values (e.g., Hyper Binding, Hyper ADMET) with user-input experimental data. When a specific R-group column is selected, the system classifies molecules into a “Structural Series” and generates group-specific graphs. This interactive interface allows users to drag and resize molecule cards on correlation graphs to visually construct a structure-activity story. This customized analysis view can be exported as a high-resolution image for immediate use in papers or reports.

### Hyper ADME/T

Hyper ADME/T can predict 19 key pharmacological and physicochemical properties including LogP, solubility, metabolic stability, CYP1A2 inhibition, CYP2C19 inhibition, CYP2C9 inhibition, CYP2D6 inhibition, CYP3A4 inhibition, CYP1A2 substrate, CYP2C19 substrate, CYP2C9 substrate, CYP2D6 substrate, CYP3A4 substrate, hERG inhibition, P-gp inhibition, PAMPA, druglikeness(9), synthesis complexity(10), and BBB penetration. From the early stages of drug development, in vitro ADME/T predictions can be used to filter molecules or inform molecular design, thereby increasing the success rate of drug development. These predicted values can be easily filtered and sorted to select desired molecules. Hyper ADME/T can perform calculations for thousands of molecules and filter them based on the predicted values.

### HyperLab AI assistant

HyperLab features an AI assistant that enhances the efficiency of drug discovery researchers. Unlike conventional LLMs, HyperLab’s AI assistant is a specialized AI agent independently developed by HITS to perform drug discovery-specific workflows. HyperLab’s AI assistant is powered by Claude 3.7 provided from AWS. It offers capabilities such as searching specialized drug discovery databases like PubChem(11), ChEMBL(7), and RCSB PDB(12), conducting web searches, coding with molecule-specific libraries, retrieving clinical trial information, visualizing graphs, and providing guidance on HyperLab usage. For example, after uploading data to HyperLab, a user can simply request “Draw a correlation between Hyper Binding score and experimental values,” and the system will generate the corresponding graph. The HyperLab AI assistant is accessible from anywhere within HyperLab and provides higher quality responses and consistency compared to general-purpose chat AIs like ChatGPT when it comes to drug discovery tasks.

Figure 8 shows an example response of HyperLab AI Assistant providing a comprehensive analysis of BI-2852, a KRAS inhibitor developed by Boehringer Ingelheim. The AI assistant explains that as of April 2025, BI-2852 has not entered clinical trials and remains primarily a research tool compound available through Boehringer Ingelheim’s opnMe platform. The response details BI-2852’s mechanism of action, including its binding to KRAS G12D, inhibition properties, and preclinical activity profile.

**Fig. 8.**
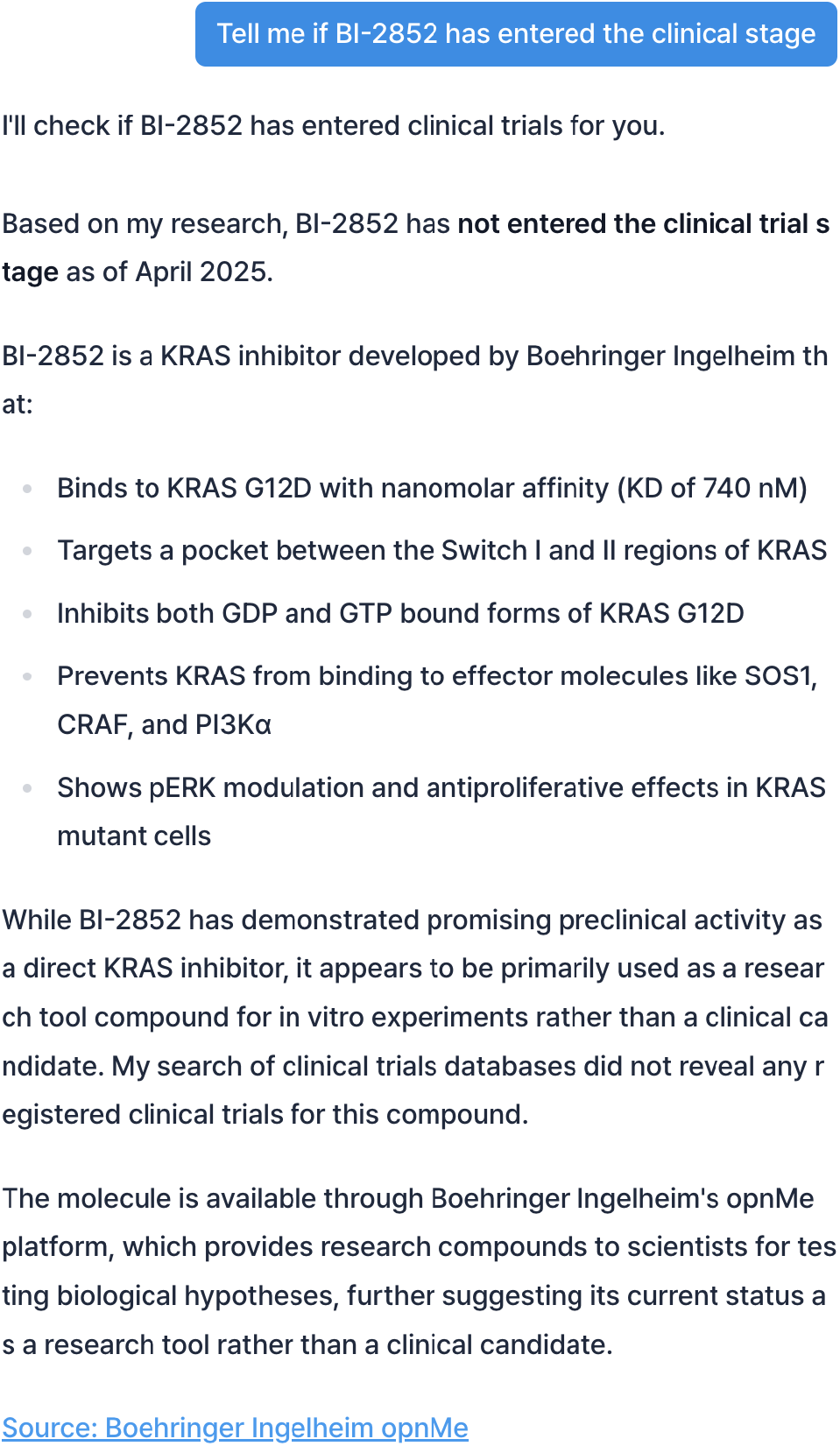
An example of how the HyperLab AI assistant would respond when a user asks about the clinical stage of BI-2852.

## Benchmark results

### Pose prediction accuracy of protein-ligand complex

To assess the pose prediction performance of various docking and structure-based modeling approaches, we conducted experiments on the PoseBuster v2(13) (PB-valid) benchmark. This benchmark is a curated dataset specifically designed to evaluate the accuracy of ligand binding pose predictions across a wide range of protein targets and chemical scaffolds, making it a rigorous testbed for assessing generalizability and robustness.

As shown in Figure 9, rigid docking methods such as Diffdock(14) and Vina(15) demonstrated limited accuracy (13% and 58%, respectively), likely due to their reliance on fixed receptor conformations and simplistic scoring functions. Chai Discovery(3) improved upon these methods (66%), while Hyper binding co-folding showed notable gains with 68% accuracy. When provided with binding site information, our method reached 77% accuracy, closely approaching the performance of AlphaFold3(16) (84%) and Boltz2(17) (78%).

**Fig. 9.**
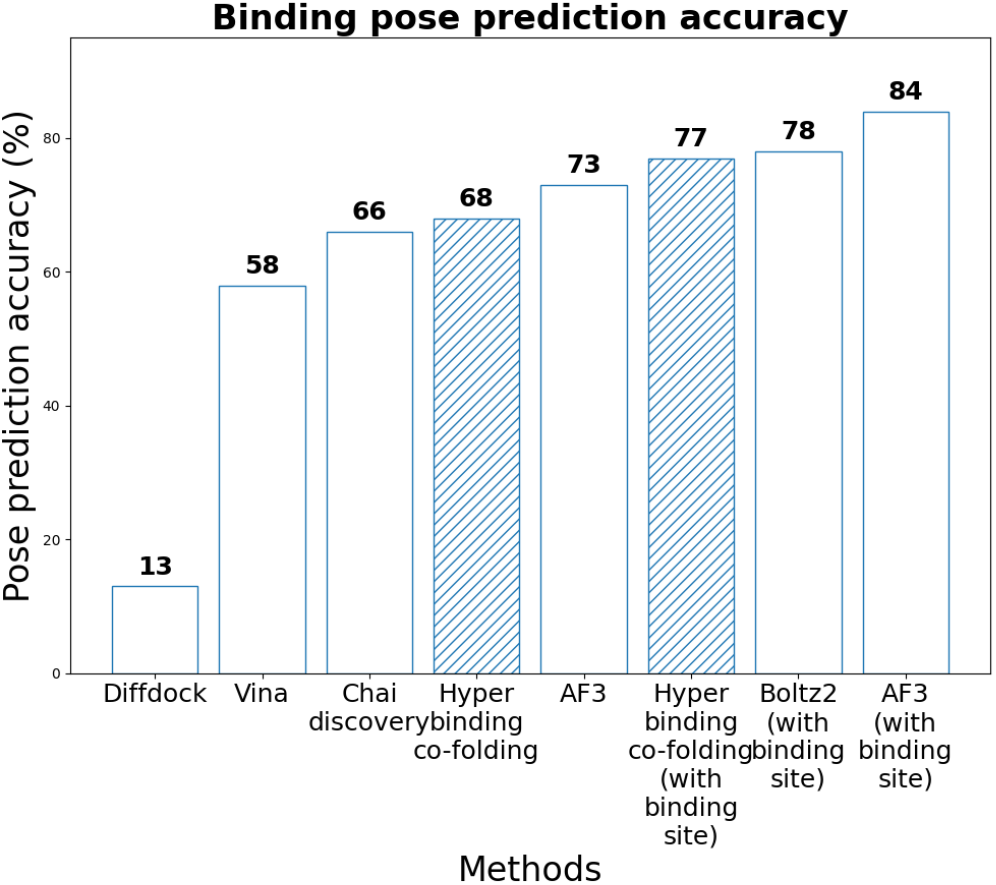
Binding pose prediction accuracy on the PoseBuster v2 (PB-valid) benchmark. Hyper binding co-folding (with binding site) achieves 77% accuracy, outperforming traditional docking methods and approaching the accuracy of AlphaFold3 (84%).

Note that unlike Vina and other docking approaches, Hyper binding co-folding does not utilize any 3D structural information of the target protein during inference. Despite this, it achieves comparable or better accuracy, highlighting the power of end-to-end complex co-folding in learning spatial constraints directly from sequence and ligand input. Importantly, our Hyper binding co-folding method offers a signif-icant advantage in inference speed. Importantly, while AlphaFold3 requires approximately 15 minutes(18) per complex with a RTX 3060 GPU, our approach completes the cofolding calculation for the same complex in just 3 minutes by using vast cloud resources. This five-fold speed advantage makes Hyper Binding co-folding a highly practical and scalable solution for virtual screening campaigns and rapid design-test-learn cycles, where computational throughput is a critical bottleneck.

### Binding affinity prediction accuracy of protein-ligand complex

Predicting the activity of derivatives sharing a common scaffold is an extremely valuable technique in hit-to-lead and lead optimization processes. However, this is a particularly challenging task because the activity differences between derivatives are often very subtle, requiring precise predictions, and small structural changes can sometimes have significant impacts on activity values. We performed benchmarking on this task using two benchmark datasets (referred to as FEP benchmark1(19) and FEP benchmark2(20)).

Figure 10 illustrates the Pearson correlation coefficient achieved by five different models on two distinct datasets, FEP dataset1 and FEP dataset2. The models evaluated are Hyper Binding, Luminet(21), Genscore(22), Glide SP(23), and Vina. On FEP dataset1, Hyper Binding demonstrated the highest performance with a Pearson correlation coefficient of 0.70. Luminet followed with a coefficient of 0.65, and Genscore achieved 0.57. Glide SP and Vina showed lower performance on this dataset, with coefficients of 0.29 and 0.25, respectively.

**Fig. 10.**
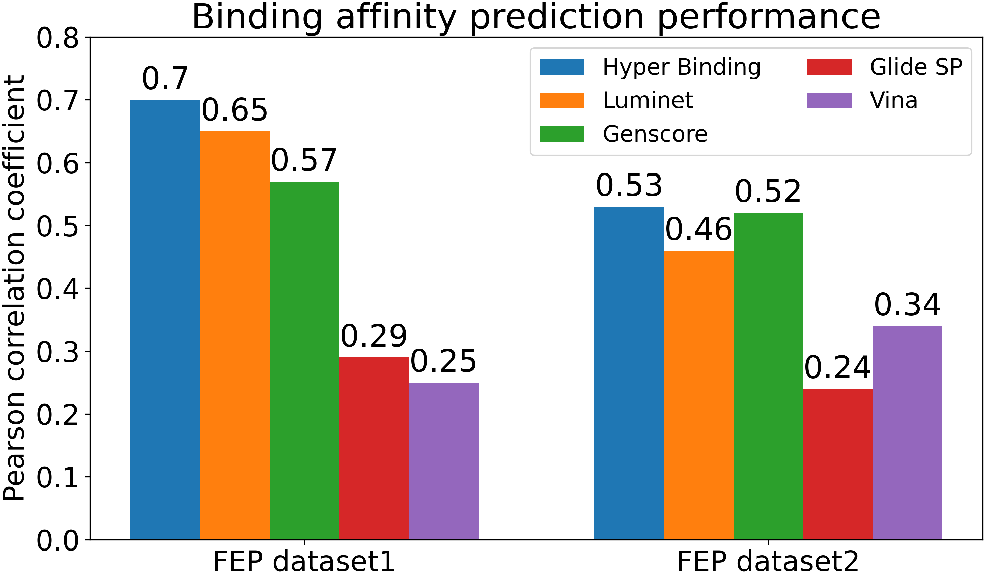
The bar chart compares the Pearson correlation coefficients of Hyper Binding against other deep learning models (Luminet, Genscore) and traditional physics-based docking methods (Glide SP, Vina). On both FEP dataset1 and FEP dataset2, Hyper Binding consistently achieved the highest correlation, demonstrating its superior accuracy in predicting binding affinity.

For FEP dataset2, Hyper Binding again achieved the highest Pearson correlation coefficient at 0.53, followed closely by Genscore at 0.52. Luminet’s performance was 0.46. Similar to FEP dataset1, docking methods such as Glide SP and Vina had the lowest scores on FEP dataset2, with coefficients of 0.24 and 0.34, respectively. Overall, Hyper Binding consistently outperformed the both deep learning models and physics based docking methods on both datasets.

### Covalent Binding affinity prediction accuracy and Screening performance

To evaluate both binding pose prediction accuracy and virtual screening enrichment of Covalent Hyper Binding, we performed a comparative benchmark analysis using a curated covalent protein–ligand dataset collected from PDBBind and related PDB entries annotated for covalent binding. The dataset spans multiple protein targets and warhead chemistries and was used as a benchmark set for assessing model robustness and generalizability. Covalent Hyper Binding was evaluated in two prediction modes, cofolding and docking. For baseline comparison, both modes were benchmarked against two widely used covalent docking tools, COV SMINA(24) and GNINA(25). To model the covalent interaction, COV SMINA imposes rigorous geometric constraints between the bonding atoms, effectively guiding the algorithm to prioritize structures in their bonded state. As shown in Figure 11, Covalent Hyper Binding (cofolding) achieved the highest pose prediction accuracy at 88.7%, clearly outperforming COV SMINA (48.4%) and GNINA (46.8%). Covalent Hyper Binding (docking) also showed competitive performance with an accuracy of 61.3%, exceeding both baseline methods. These results indicate that both HyperLab modes provide stronger pose prediction performance than the conventional baseline tools evaluated here. To assess virtual screening performance, the enrichment factor at 10% (EF@10%) was calculated.This metric measures how strongly a method concentrates actives near the top of the ranking relative to random selection. EF@10% was evaluated for Mpro and KRAS using a common library of 300 compounds containing 3 actives and 297 decoys (top 10% = 30 compounds). All methods were tested on the same candidate pool under identical conditions. Covalent Hyper Binding and COV SMINA poses were ranked with the HyperLab Covalent Hyper Binding score because COV SMINA does not provide a dedicated covalent scoring function. GNINA poses were ranked with the GNINA score, which is not specific to covalent systems. For Mpro, Covalent Hyper Binding (docking) achieved the highest EF@10% at 6.56, while the other methods showed no enrichment in the top-ranked 10% fraction. For KRAS, Covalent Hyper Binding (docking) again showed the strongest enrichment at 9.97, followed by COV SMINA at 3.32, while GNINA showed no enrichment. Overall, Covalent Hyper Binding workflow showed the strongest prioritization performance in this screening setting.

**Fig. 11.**
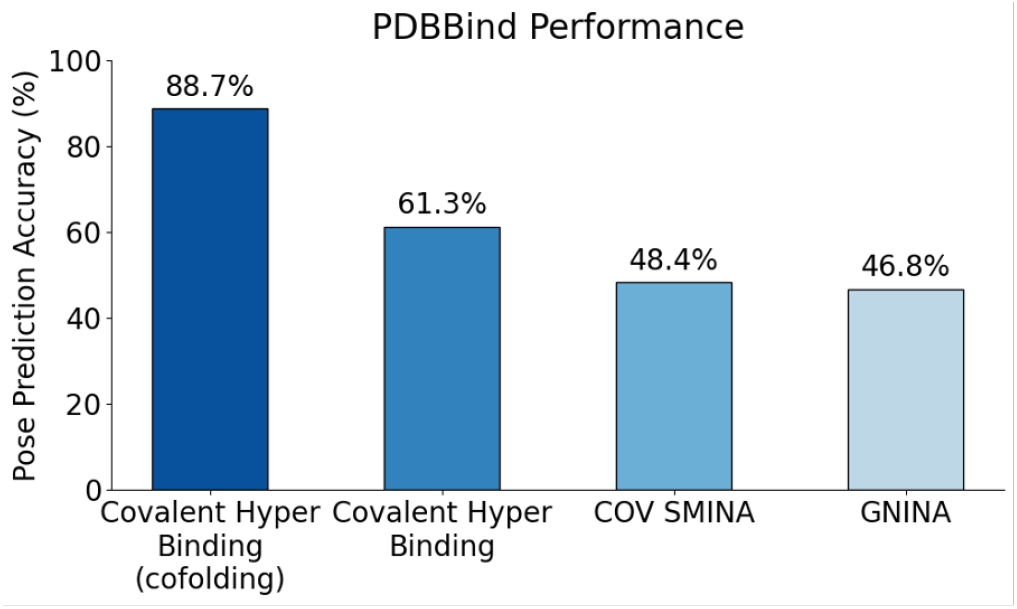
The firgure presents a benchmark comparison using the covalent data of PDBBind dataset, demonstrating that Covalent Hyper Binding achieves significantly higher pose prediction accuracy than traditional docking tools like COV SMINA and GNINA.

**Table 2.** Comparison of covalent docking performance between Covalent Hyper Binding and baseline methods (COV SMINA, GNINA) for Mpro and KRAS targets.

| Method | Mpro | KRAS |
| --- | --- | --- |
| Covalent Hyper Binding | 6.56 | 9.97 |
| COV SMINA | 0.00 | 3.32 |
| GNINA | 0.00 | 0.00 |

## Case study: AI-enabled screening and design with HyperLab

HyperLab was created to compress the early phases of drug discovery by combining ultra-fast virtual screening (“Hyper Screening”) with an AI-guided compound-design module (“Hyper Design”). To evaluate the platform’s technical performance, we conducted an internal validation study using a pipeline developed in-house. In the Hyper Screening stage, we selected the PDB structure of the target protein and performed Hyper Binding calculations against a diverse compound library. Based on binding scores, we directly selected 52 top-ranked compounds for in vitro testing. The entire virtual screening process was completed within 24 hours. In vitro assay results (Figure 12) identified five compounds showing >40% inhibition, corresponding to a hit rate of approximately 9%. The IC_50_ values of the identified compounds, including the reference compound H00112 (IC_50_=1319 nM), ranged from 70 to 600 nM.

**Fig. 12.**
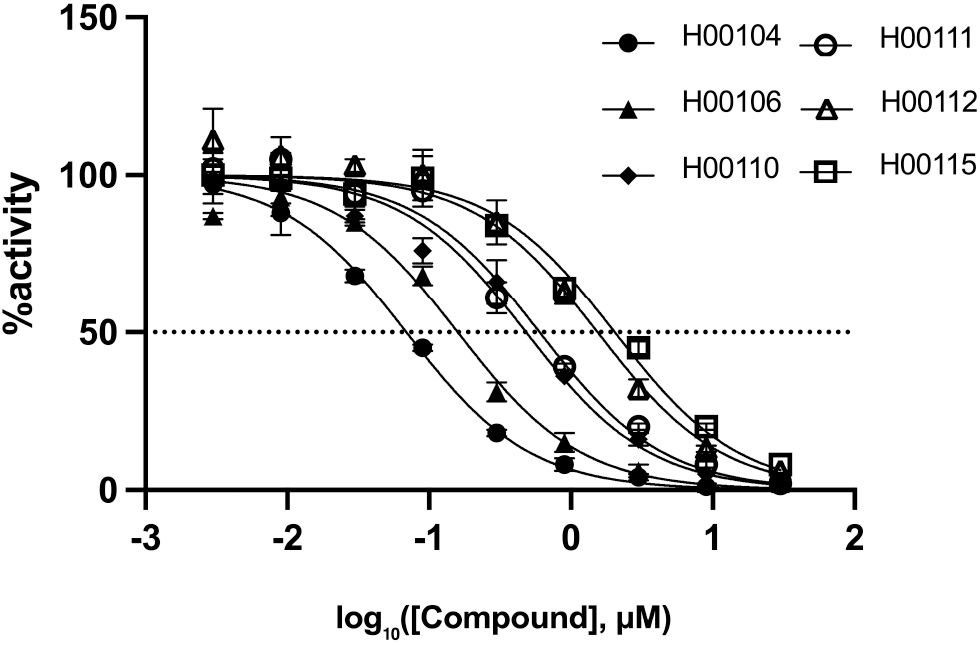
Hit validation of top-ranked compounds from Hyper Screening. Dose–response curves for the six hit compounds identified in, measured in a kinase inhibition assay. Curves were fitted using a standard sigmoid model to determine IC_50_ values. All data are expressed as mean ± SD from duplicate experiments, relative to the DMSO control. Assays were performed by Eurofins Discovery using the KinaseProfiler LeadHunter radiometric platform.

Using the Hyper Design, we conducted a part-by-part replacement of the reference compound to generate structurally novel derivatives. Compound selection was guided by multiple criteria, including improvement in predicted binding scores, retention of key molecular interactions with the target protein, and structural novelty with a particular emphasis on avoiding patent-protected motifs of the reference compound. Among the 14 candidate compounds selected through this process, 10 (71%) were assessed to be synthetically accessible based on human inspection. Five of these were synthesized early and subsequently subjected to in vitro assays. The three compounds exhibited over 75% inhibition at a concentration of 1 *µ*M, with IC_50_ values ranging from 200 to 400 nM. These results are shown in Figure Fig. 13.

**Fig. 13.**
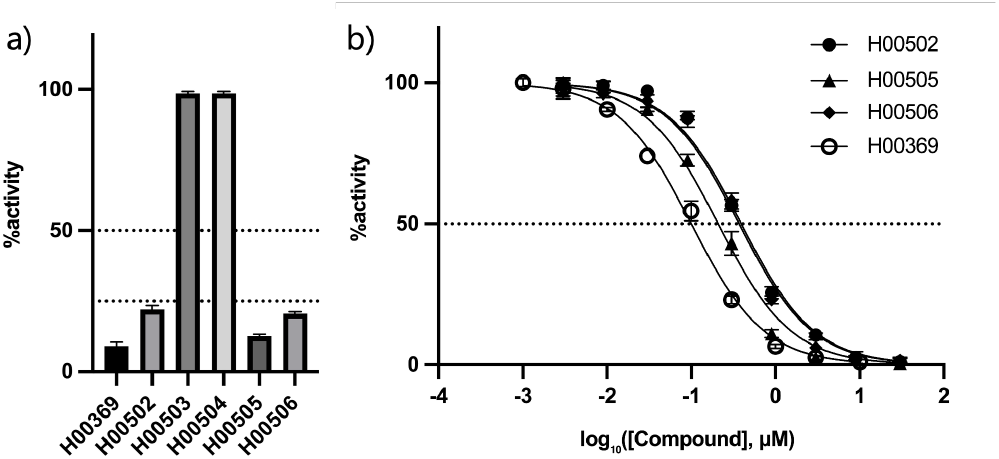
Inhibitory activity of synthesized compounds against the target protein. Single-dose assay at 1 µM showing the percentage activity of the target protein relative to the DMSO control (set as 100%). H00369 is the reference compound, and all others are newly synthesized derivatives. b) Dose–response curves of selected compounds (H00502, H00505, H00506, and H00369) plotted as percentage activity versus log_10_ [compound] (*µ*M). All data are expressed as mean ± SD from duplicate experiments, relative to the DMSO control. Assays were performed by BPS Bioscience

H00505 emerged as the most potent compound among those tested, exhibiting inhibitory activity comparable to that of the reference compound H00369. This finding prompted further mechanistic studies to evaluate its engagement with the relevant signaling pathway. In the pathway-specific assay, measured by the ARE luciferase reporter system (Figure Fig. 14), H00505 demonstrated activity equal to or greater than that of the reference compound, highlighting its potential as a promising lead candidate for further optimization.

**Fig. 14.**
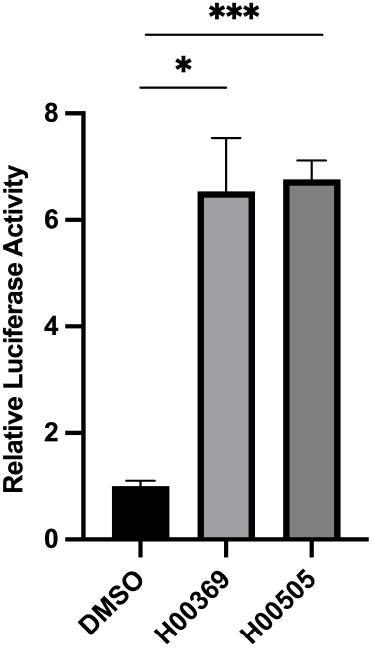
The ARE transcriptional activity of H00505. ARE–luciferase–HepG2 cells were treated for 18 h with 10*µ*M of H00505 and the reference compound, and luciferase activity was measured using the ONE-Step™ Luciferase Assay System (BPS Bioscience). H00505 induced a significantly greater fold increase in ARE activity compared to both the DMSO control and the reference compound, indicating enhanced activation of the antioxidant response pathway. Data are presented as mean ± SD from triplicate experiments, relative to the DMSO control. p values were determined using an unpaired two-tailed t-test. p<0.05 (*), p<0.001 (***).

Taken together, these results demonstrate that Hyper-Lab efficiently identifies novel hit compounds and converts in-silico designs into experimentally validated leads with nanomolar potency, supporting its application as a scalable and cost-effective platform for target-based drug discovery.

Conventional high-throughput screening (HTS) campaigns typically interrogate ≥105 compounds in single-concentration assays, with subsequent confirmatory titrations to obtain IC_50_ values, and report primary hit rates in the low single-digit percent range. In contrast, Hyper Screening identified 6 actives (11.5%) from only 52 candidates, while Hyper Design yielded 3 actives out of 5 synthesized (60%), thereby introducing structural differentiation without loss of potency. Assuming standard assay workflows (duplicate single-point and 10-point duplicate titrations), this corresponds to a reduction from 200,000 wells in a representative HTS to 100 wells in our pipelines. These results demonstrate the efficiency gains achievable by computational triage and AI-guided motif design in reducing both time and cost while maintaining pharmacological relevance.

## Conclusion

HyperLab represents a significant advancement in AI-driven drug discovery platforms, addressing the critical need for accessible computational tools for drug discovery researchers. By combining state-of-the-art AI models with an intuitive user interface, HyperLab democratizes access to sophisticated drug discovery capabilities that were previously available only to computational specialists. The platform’s comprehensive suite of tools—Hyper Binding, Hyper Design, Hyper Screening, and Hyper ADME/T—covers the entire spectrum of structure-based drug discovery needs, from initial hit identification to lead optimization.

Our benchmark results demonstrate HyperLab’s superior performance compared to conventional methods, with Hyper Binding achieving 77% accuracy in binding pose prediction and consistently outperforming both deep learning models and physics-based docking methods in binding affinity prediction across multiple datasets. These performance metrics, combined with the platform’s user-friendly interface and specialized AI assistant, position HyperLab as a transformative tool for accelerating drug discovery processes.

The practical power of this integrated platform was validated through an internal case study, which progressed from initial high-throughput screening to the experimental confirmation of a novel lead compound with nanomolar potency (IC_50_ values of 200-400 nM). This achievement demonstrates HyperLab’s real-world capability to dramatically compress discovery timelines and efficiently convert insilico designs into experimentally validated leads. As variety of AI methods continue to play an increasingly vital role in pharmaceutical research, HyperLab stands poised to significantly reduce the time, cost, and expertise barriers associated with modern drug discovery efforts.

## Supporting information

all latex file

## Revision History

**2026-04** Covalent feature release update.

**2026-01** SAR feature release update.

**2025-09** Initial release of the HyperLab platform documentation.

## Notes

### Competing Interest Statement

This is the publication from the profit organization and it is about their product.

### Summary of Updates

The following updates regarding the HyperLab covalent module have been incorporated into the latest bioRxiv submission: The updated manuscript introduces HyperLab's new covalent module, which now provides a comprehensive and streamlined workflow for binding prediction, molecular design, and virtual screening. Central to this update is the introduction of a dual-mode system: an Auto-Recommendation mode that enables automatic warhead-residue matching, and a Manual Definition mode for highly customized configurations. By leveraging advanced co-folding technology, the platform can now predict complex structures directly from protein sequences, significantly enhancing its utility in cases where pre-resolved structures are unavailable. The submission also highlights a major leap in performance benchmarks. Covalent Hyper Binding (co-folding) achieved a remarkable 88.7% pose prediction accuracy, which represents a substantial improvement over traditional industry tools like COV SMINA (48.4%) and GNINA (46.8%). The module's superior screening power is further evidenced by its impressive Enrichment Factors (EF@10%), reaching 9.97 for KRAS and 6.56 for Mpro. Finally, the revised text emphasizes the module's full integration with SAR analysis and Hyper ADME/T. This allows researchers to evaluate both the reactant and covalent product states within a single, unified interface, ensuring a more holistic and efficient approach to covalent drug discovery.

## Bibliography

1. Mihaly Varadi, Damian Bertoni, Paulyna Magana, Urmila Paramval, Ivanna Pidruchna, Malarvizhi Radhakrishnan, Maxim Tsenkov, Sreenath Nair, Milot Mirdita, Jingi Yeo, Oleg Kovalevskiy, Kathryn Tunyasuvunakool, Agata Laydon, Augustin Žídek, Hamish Tomlinson, Dhavanthi Hariharan, Josh Abrahamson, Tim Green, John Jumper, Ewan Birney, Martin Steinegger, Demis Hassabis, and Sameer Velankar. Alphafold protein structure database in 2024: providing structure coverage for over 214 million protein sequences. Nucleic Acids Research, 52:D368–D375, 1 2024. ISSN 0305-1048. doi: 10.1093/nar/gkad1011.

2. Seokhyun Moon, Sang-Yeon Hwang, Jaechang Lim, and Woo Youn Kim. Pignet2: a versatile deep learning-based protein–ligand interaction prediction model for binding affinity scoring and virtual screening. Digital Discovery, 3:287–299, 2024. ISSN 2635-098X. doi: 10.1039/D3DD00149K.

3. Jacques Boitreaud, Jack Dent, Matthew McPartlon, Joshua Meier, Vinicius Reis, Alex Rogozhnikov, and Kevin Wu. Chai-1: Decoding the molecular interactions of life, 10 2024.

4. Jiankun Lyu, Sheng Wang, Trent E. Balius, Isha Singh, Anat Levit, Yurii S. Moroz, Matthew J. O’Meara, Tao Che, Enkhjargal Algaa, Kateryna Tolmachova, Andrey A. Tolmachev, Brian K. Shoichet, Bryan L. Roth, and John J. Irwin. Ultra-large library docking for discovering new chemotypes. Nature, 566:224–229, 2 2019. ISSN 0028-0836. doi: 10.1038/s41586-019-0917-9.

5. Emmanuel Bengio, Moksh Jain, Maksym Korablyov, Doina Precup, and Yoshua Bengio. Flow network based generative models for non-iterative diverse candidate generation. 11 2021.

6. Seonghwan Seo, Minsu Kim, Tony Shen, Martin Ester, Jinkyoo Park, Sungsoo Ahn, and Woo Youn Kim. Generative flows on synthetic pathway for drug design. 3 2025.

7. Barbara Zdrazil, Eloy Felix, Fiona Hunter, Emma J Manners, James Blackshaw, Sybilla Corbett, Marleen de Veij, Harris Ioannidis, David Mendez Lopez, Juan F Mosquera, Maria Paula Magarinos, Nicolas Bosc, Ricardo Arcila, Tevfik Kizilören, Anna Gaulton, A Patrícia Bento, Melissa F Adasme, Peter Monecke, Gregory A Landrum, and Andrew R Leach. The chembl database in 2023: a drug discovery platform spanning multiple bioactivity data types and time periods. Nucleic Acids Research, 52:D1180–D1192, 1 2024. ISSN 0305-1048. doi: 10.1093/nar/gkad1004.

8. Michael Deininger, Elisabeth Buchdunger, and Brian J. Druker. The development of imatinib as a therapeutic agent for chronic myeloid leukemia. Blood, 105:2640–2653, 4 2005. ISSN 0006-4971. doi: 10.1182/blood-2004-08-3097.

9. Kyunghoon Lee, Jinho Jang, Seonghwan Seo, Jaechang Lim, and Woo Youn Kim. Druglikeness scoring based on unsupervised learning. Chemical Science, 13:554–565, 2022. ISSN 2041-6520. doi: 10.1039/D1SC05248A.

10. Hyeongwoo Kim, Kyunghoon Lee, Chansu Kim, Jaechang Lim, and Woo Youn Kim. Dfrscore: Deep learning-based scoring of synthetic complexity with drug-focused retrosynthetic analysis for high-throughput virtual screening. Journal of Chemical Information and Modeling, 64:2432–2444, 4 2024. ISSN 1549-9596. doi: 10.1021/acs.jcim.3c01134.

11. Sunghwan Kim, Jie Chen, Tiejun Cheng, Asta Gindulyte, Jia He, Siqian He, Qingliang Li, Benjamin A Shoemaker, Paul A Thiessen, Bo Yu, Leonid Zaslavsky, Jian Zhang, and Evan E Bolton. Pubchem 2025 update. Nucleic Acids Research, 53:D1516–D1525, 1 2025. ISSN 0305-1048. doi: 10.1093/nar/gkae1059.

12. Christine Zardecki, Shuchismita Dutta, David S. Goodsell, Robert Lowe, Maria Voigt, and Stephen K. Burley. Pdb -101: Educational resources supporting molecular explorations through biology and medicine. Protein Science, 31:129–140, 1 2022. ISSN 0961-8368. doi: 10.1002/pro.4200.

13. Martin Buttenschoen, Garrett M. Morris, and Charlotte M. Deane. Posebusters: Ai-based docking methods fail to generate physically valid poses or generalise to novel sequences. Chemical Science, 15:3130–3139, 2024. ISSN 2041-6520. doi: 10.1039/D3SC04185A.

14. Gabriele Corso, Hannes Stärk, Bowen Jing, Regina Barzilay, and Tommi Jaakkola. Diffdock: Diffusion steps, twists, and turns for molecular docking. 2 2023.

15. Jerome Eberhardt, Diogo Santos-Martins, Andreas F. Tillack, and Stefano Forli. Autodock vina 1.2.0: New docking methods, expanded force field, and python bindings. Journal of Chemical Information and Modeling, 61:3891–3898, 8 2021. ISSN 1549-9596. doi: 10.1021/acs.jcim.1c00203.

16. Josh Abramson, Jonas Adler, Jack Dunger, Richard Evans, Tim Green, Alexander Pritzel, Olaf Ronneberger, Lindsay Willmore, Andrew J. Ballard, Joshua Bambrick, Sebastian W. Bodenstein, David A. Evans, Chia-Chun Hung, Michael O’Neill, David Reiman, Kathryn Tunyasuvunakool, Zachary Wu, Akvilė Žemgulytė, Eirini Arvaniti, Charles Beattie, Ottavia Bertolli, Alex Bridgland, Alexey Cherepanov, Miles Congreve, Alexander I. Cowen-Rivers, Andrew Cowie, Michael Figurnov, Fabian B. Fuchs, Hannah Gladman, Rishub Jain, Yousuf A. Khan, Caroline M. R. Low, Kuba Perlin, Anna Potapenko, Pascal Savy, Sukhdeep Singh, Adrian Stecula, Ashok Thillaisundaram, Catherine Tong, Sergei Yakneen, Ellen D. Zhong, Michal Zielinski, Augustin Žídek, Victor Bapst, Pushmeet Kohli, Max Jaderberg, Demis Hassabis, and John M. Jumper. Accurate structure prediction of biomolecular interactions with alphafold 3. Nature, 630:493–500, 6 2024. ISSN 0028-0836. doi:10.1038/s41586-024-07487-w.

17. Saro Passaro, Gabriele Corso, Jeremy Wohlwend, Mateo Reveiz, Stephan Thaler, Vignesh Ram Somnath, Noah Getz, Tally Portnoi, Julien Roy, Hannes Stark, David Kwabi-Addo, Dominique Beaini, Tommi Jaakkola, and Regina Barzilay. Boltz-2: Towards accurate and efficient binding affinity prediction, 6 2025.

18. Taras Voitsitskyi, Ihor Koleiev, Roman Stratiichuk, Oleksandr Kot, Roman Kyrylenko, Illia Savchenko, Vladyslav Husak, Semen Yesylevskyy, Sergii Starosyla, and Alan Nafiiev. Artidock: accurate machine learning approach to protein-ligand docking optimized for high-throughput virtual screening, 3 2024.

19. Lingle Wang, Yujie Wu, Yuqing Deng, Byungchan Kim, Levi Pierce, Goran Krilov, Dmitry Lupyan, Shaughnessy Robinson, Markus K. Dahlgren, Jeremy Greenwood, Donna L. Romero, Craig Masse, Jennifer L. Knight, Thomas Steinbrecher, Thijs Beuming, Wolfgang Damm, Ed Harder, Woody Sherman, Mark Brewer, Ron Wester, Mark Murcko, Leah Frye, Ramy Farid, Teng Lin, David L. Mobley, William L. Jorgensen, Bruce J. Berne, Richard A. Friesner, and Robert Abel. Accurate and reliable prediction of relative ligand binding potency in prospective drug discovery by way of a modern free-energy calculation protocol and force field. Journal of the American Chemical Society, 137:2695–2703, 2 2015. ISSN 0002-7863. doi: 10.1021/ja512751q.

20. Christina E. M. Schindler, Hannah Baumann, Andreas Blum, Dietrich Böse, Hans-Peter Buchstaller, Lars Burgdorf, Daniel Cappel, Eugene Chekler, Paul Czodrowski, Dieter Dorsch, Merveille K. I. Eguida, Bruce Follows, Thomas Fuchß, Ulrich Grädler, Jakub Gunera, Theresa Johnson, Catherine Jorand Lebrun, Srinivasa Karra, Markus Klein, Tim Knehans, Lisa Koetzner, Mireille Krier, Matthias Leiendecker, Birgitta Leuthner, Liwei Li, Igor Mochalkin, Djordje Musil, Constantin Neagu, Friedrich Rippmann, Kai Schiemann, Robert Schulz, Thomas Steinbrecher, Eva-Maria Tanzer, Andrea Unzue Lopez, Ariele Viacava Follis, Ansgar Wegener, and Daniel Kuhn. Large-scale assessment of binding free energy calculations in active drug discovery projects. Journal of Chemical Information and Modeling, 60:5457–5474, 11 2020. ISSN 1549-9596. doi: 10.1021/acs.jcim.0c00900.

21. Qun Su, Jike Wang, Qiaolin Gou, Renling Hu, Linlong Jiang, Hui Zhang, Tianyue Wang, Yifei Liu, Chao Shen, Yu Kang, Chang-Yu Hsieh, and Tingjun Hou. Robust protein–ligand interaction modeling through integrating physical laws and geometric knowledge for absolute binding free energy calculation. Chemical Science, 16:5043–5057, 2025. ISSN 2041-6520. doi: 10.1039/D4SC07405J.

22. Chao Shen, Xujun Zhang, Chang-Yu Hsieh, Yafeng Deng, Dong Wang, Lei Xu, Jian Wu, Dan Li, Yu Kang, Tingjun Hou, and Peichen Pan. A generalized protein–ligand scoring framework with balanced scoring, docking, ranking and screening powers. Chemical Science, 14:8129–8146, 2023. ISSN 2041-6520. doi: 10.1039/D3SC02044D.

23. Richard A. Friesner, Jay L. Banks, Robert B. Murphy, Thomas A. Halgren, Jasna J. Klicic, Daniel T. Mainz, Matthew P. Repasky, Eric H. Knoll, Mee Shelley, Jason K. Perry, David E. Shaw, Perry Francis, and Peter S. Shenkin. Glide: a new approach for rapid, accurate docking and scoring. 1. method and assessment of docking accuracy. Journal of Medicinal Chemistry, 47:1739–1749, 3 2004. ISSN 0022-2623. doi: 10.1021/jm0306430.

24. David Ryan Koes, Matthew P. Baumgartner, and Carlos J. Camacho. Lessons learned in empirical scoring with smina from the csar 2011 benchmarking exercise. Journal of Chemical Information and Modeling, 53:1893–1904, 8 2013. ISSN 1549-9596. doi: 10.1021/ci300604z.

25. Andrew T. McNutt, Yanjing Li, Rocco Meli, Rishal Aggarwal, and David Ryan Koes. Gnina 1.3: the next increment in molecular docking with deep learning. Journal of Cheminformatics, 17:28, 3 2025. ISSN 1758-2946. doi: 10.1186/s13321-025-00973-x.

