## Supplementary material for "Accelerating Drug Discovery with HyperLab: An Easy-to-Use AI-Driven Platform": all latex file: Article.tex

\documentclass[times, twoside, watermark]{zHenriquesLab-StyleBioRxiv}
\usepackage[utf8]{inputenc}
\usepackage{blindtext}
% \usepackage{hyperref}
\usepackage{booktabs}
\usepackage{booktabs}  % 가로선 굵기 조절 (\toprule, \midrule, \bottomrule)
\usepackage{tabularx}  % 표 너비 자동 조절
\usepackage{ragged2e}  % 텍스트 정렬 개선
%\usepackage{cleveref}

% Please give the surname of the lead author for the running footer
\leadauthor{Lim} 

\begin{document}

\title{Accelerating Drug Discovery with HyperLab: An Easy-to-Use AI-Driven Platform}
\shorttitle{HyperLab technical report}

% Use letters for affiliations, numbers to show equal authorship (if applicable) and to indicate the corresponding author
\author[1]{HyperLab team}
}

\affil[1]{HITS, 8F, 28, Teheran-ro 4-gil, Gangnam-gu, Seoul, Republic of Korea}

    \maketitle

%TC:break Abstract
%the command above serves to have a word count for the abstract
\begin{abstract}

HyperLab, developed by HITS, is a web-based, AI-driven drug discovery platform designed to increase research efficiency for experimental drug discovery researchers. The platform features an intuitive user interface and experience (UI/UX), enabling researchers without specialized expertise in AI or computational methods to readily generate essential discovery outcomes. By employing a Structure-Based Drug Discovery (SBDD) methodology, HyperLab streamlines the complete discovery workflow. Its core functionalities include the prediction of ligand-protein structures and binding activities (Hyper Binding), protein structure-based molecular optimization (Hyper Design), structure-activity-relationship (SAR) analysis, screening of extensive chemical libraries ranging from one million to seven trillion compounds (Hyper Screening and Hyper Screening X), prediction of ADMET properties (Hyper ADME/T), and an AI-driven assistant designed to boost researcher productivity and efficiency.

In benchmark evaluations, Hyper Binding demonstrated 77\% accuracy for binding pose prediction on the PoseBuster v2 benchmark, surpassing conventional docking techniques and closely approaching the 84\% accuracy of AlphaFold3, while offering a considerable advantage in computational speed. Furthermore, for binding affinity prediction, HyperBinding showed superior performance over both deep learning and physics-based docking models on two distinct FEP datasets, achieving Pearson correlation coefficients of 0.70 and 0.53, respectively.

For experimental validation, an internal study utilized Hyper Screening to identify top-ranked compounds without without any post-analysis or visual inspection by human experts. The screening process was completed in 24 hours. Through experimental validation of top-ranked compounds, five compounds with IC$_{50}$ values ranging from 70 to 600 nM were identified. Hyper Design was employed to generate novel derivatives with improved predicted binding scores and structural novelty. Of five synthesized compounds, \textit{in vitro} assays confirmed that three demonstrated over 75\% inhibition at a 1~\textmu M concentration, with IC\textsubscript{50} values in the 200 to 400~nM range. Notably, one compound exhibited activity comparable or superior to the reference compound. HyperLab is therefore positioned to substantially lower the barriers of time, cost, and specialized expertise inherent in modern drug discovery initiatives.
\end{abstract}

%TC:break main
%the command above serves to have a word count for the abstract

\begin{keywords}
AI drug discovery platform | AI | drug discovery | HyperLab
\end{keywords}

\begin{corrauthor}
%\texttt{jaechang{@}hits.ai}
jaechang\at hits.ai
\end{corrauthor}

\section*{Introduction}
HyperLab is an web-based AI-driven drug discovery platform developed by HITS for pharmaceutical researchers. HyperLab is specifically designed for use by all pharmaceutical researchers in drug development, enabling them to leverage its capabilities without needing specialized expertise in computer aided drug discovery. Users can obtain necessary results easily through an intuitive UI/UX, without requiring specialized knowledge or experience in AI or CADD. Key features of HyperLab are predictions of protein-ligand structures and activities, protein structure-based molecular optimization, chemical library screening (from 1 million to 7 trillion compounds), ADME/T prediction, and AI assistant that enhances researcher productivity and efficiency. This technical report explains HyperLab's main functionalities, underlying technologies, and predictive performance.

\section*{Overview of HyperLab key functionalities}
HyperLab has developed state-of-the-art protein structure-based AI models integrated with an intuitive user interface. The primary objective of HyperLab is to facilitate the expeditious identification and optimization of compounds with optimal inhibitory activity against target proteins and variety of pharmacological properties.

HyperLab is an integrated AI platform that streamlines the entire drug discovery pipeline. The workflow can begin with Hyper Binding for accurate protein-ligand binding prediction and affinity assessment. HyperLab also provides optimized worflow for hit identification through Hyper Screening allowing virtual screening of millions of compounds and Hyper Screening X for multi-parameter optimization across trillions of molecules in ultra-large virtual chemical spaces. The platform then transitions to lead optimization using Hyper ADME/T for comprehensive ADMET property prediction and Hyper Design for structure-based molecular optimization to enhance binding affinity while maintaining drug-like properties. To further empower the Hit-to-Lead and Lead Optimization stages, HyperLab incorporates a specialized Structure-Activity Relationship (SAR) analysis module. This integrated approach, supported by an AI assistant throughout the process, significantly reduces the time and resources required for early-stage drug discovery by providing seamless transitions between traditionally fragmented workflows within a single, cohesive environment.

For targets with experimentally determined three-dimensional structures, researchers can initialize the protein structure by specifying a PDB identifier. The proprietary database on the platform enables automatic identification of binding sites and configuration of appropriate binding boxes. Researchers retain the flexibility to manually adjust the locations and dimensions of the binding site as needed. Additionally, the platform supports direct upload of custom PDB files for protein structure initialization. In cases where experimental structures are unavailable, the system can utilize AlphaFold-predicted structures\cite{Varadi2024} by referencing UniProt identifiers.

Furthermore, HyperLab has implemented advanced Co-folding technology that enables prediction of protein-ligand complex binding conformations directly from protein sequences. This Co-folding methodology significantly streamlines the protein registration workflow by eliminating the need to select specific PDB structures. Researchers need only input the protein name to retrieve sequence data, binding site information, and post-translational modification details from the comprehensive pre-constructed database. Following protein registration, users can modify structural parameters and incorporate multiple proteins to predict binding interactions between protein complexes and ligands. The platform also supports direct input of custom protein sequences independently of the database.

\begin{figure}%[tbhp]
\centering
\includegraphics[width=1.0\linewidth]{Figures/overview.png}
\caption{The integrated pipeline demonstrates the comprehensive drug discovery process from hit identification to lead optimization. The workflow progresses through three main phases: (1) hit and target Characterization using Hyper Binding for protein-ligand binding prediction, (2) hit Discovery through virtual screening of millions of compounds and advanced Hyper Screening X for multi-parameter optimization across trillions of molecules, and (3) hit-to-lead and lead optimization employing Hyper ADME/T for ADMET property prediction and Hyper Design for structure-based molecular optimization. The AI assistant provides continuous support throughout the entire discovery process.}
\label{fig:overview}
\end{figure}

\subsection*{Hyper Binding}
Hyper Binding represents an advanced key function of HyperLab for the prediction of protein-ligand interactions and binding conformations. Hyper binding enhanced generalization performance through the use of a physics-informed deep learning model\cite{Moon2024}. The system accepts varierty of protein structure input format, including PDB identifier registration, user-uploaded structures, and AlphaFold-predicted conformations as explained in the previous parts. In Hyper Binding, the algorithm explores potential ligand-binding poses within user-defined binding sites while maintaining the protein structure as a rigid entity. Additionally, for cases where only protein sequence information is available, Hyper Binding implements a co-folding approach\cite{Boitreaud2024} that simultaneously predicts both the protein structure and ligand binding conformation.

The platform provides comprehensive visualization capabilities, enabling two-dimensional and three-dimensional analysis of predicted binding modes. These structural insights facilitate the mechanistic understanding of protein-ligand interactions and inform rational structure modification strategies to enhance binding affinity. Furthermore, Hyper Binding supports selectivity profiling through comparative binding predictions across multiple protein targets.

Also, Hyper Binding can be utilized to determine the bioassay priority. After uploading a list of candidate molecules designed by medicinal chemists, binding scores can be predicted to establish bioassay priorities. Through cloud-based parallelization, hundreds to thousands of molecules can be rapidly processed, significantly accelerating the computational workflow.

Figure \labelcref{fig:hyperbinding}. shows the binding mode analysis of Pirtobrutinib (LOXO-305), a selective and non-covalent Bruton's tyrosine kinase (BTK) inhibitor used in cancer treatment in complex with BTK produced by Hyper Binding. 
Figure \labelcref{fig:hyperbinding}-a) displays the three-dimensional representation of Pirtobrutinib bound to the ATP-binding pocket of BTK (PDB ID: 8FLL). The protein is shown as a blue ribbon structure with key interacting residues displayed as sticks. The predicted ligand is represented in purple stick format, highlighting its alignment with the X-ray structure represented in cyan stick format. The figure shows the orientation of the ligand within the binding pocket and key interactions with surrounding amino acid residues. Figure \labelcref{fig:hyperbinding}-b) shows the two-dimensional interaction diagram of Pirtobrutinib with BTK residues calculated using Hyper Binding. The diagram illustrates various interaction types including hydrogen bonds (pink dashed lines), cation-π interactions (yellow dashed lines), and π-π stacking (light pink). Key residues involved in binding include MET477, LEU528, CYS481, and LYS430. The trifluoromethyl pyrazole group, central phenyl ring, and methoxy-substituted benzamide moiety of Pirtobrutinib form multiple favorable interactions within the binding pocket, explaining its potent and selective inhibition of BTK.

\begin{figure}%[tbhp]
\centering
\includegraphics[width=1.0\linewidth]{Figures/hyperbinding.png}
\caption{Visualization of Pirtobrutinib (LOXO-305) in complex with Bruton's tyrosine kinase: (a three-dimensional structural representation and (b two-dimensional interaction diagram. }
\label{fig:hyperbinding}
\end{figure}

\subsection*{Hyper Screening}
Hyper Screening is a virtual screening function designed to discover initial active compounds in the early-stage of drug discovery. It performs Hyper Binding calculations on all molecules in a library against a specified protein, then provides the top 500 molecules with the best binding scores. Five pre-processed libraries are available for specific purposes. When Hyper Screening is performed using the provided libraries, compound information and purchase links are provided to enable users to acquire these substances. Users can also register their own libraries to perform screening. Once a library is registered, it can be used for screening against different targets. For the selected top 500 molecules, users can perform 2D/3D interaction analysis and optimize molecular structures using Hyper Design, just as with regular Hyper Binding calculations.

The Table \labelcref{tab:hyperscreening} summarizes the five primary molecule libraries available in Hyper Screening. The Fragment library contains 500,000 molecules selected from the Diverse library that meet specific physicochemical criteria (MW < 300, LogP < 3.0, < 4 hydrogen bond donors). The comprehensive Diverse library includes 1 million diverse compounds filtered according to HITS's proprietary criteria, including Lipinski rule compliance and clustering based on in-stock availability. The FDA-approved library comprises 1,100 marketed drugs, while the Natural product-like fragments library contains 4,200 molecules representing 1,000 structural motifs derived from natural products. The Kinase-focused library of 65,000 compounds includes bioisosteric derivatives of allosteric kinase inhibitors and hinge binder molecules specifically designed for kinase-targeted drug discovery project.

\begin{table}[htbp]
    \centering
    \caption{Overview of Hyper Screening libraries}
    \renewcommand{\arraystretch}{1.4} % 행 높이를 넉넉하게
    % 컬럼 설정 설명:
    % 1열: 1.75cm, 왼쪽 정렬
    % 2열: 1.3cm, 오른쪽 정렬 (숫자 보기 좋게)
    % 3열: 4.0cm, 왼쪽 정렬
    \begin{tabular}{ >{\raggedright\arraybackslash}p{1.75cm} >{\raggedleft\arraybackslash}p{1.3cm} >{\raggedright\arraybackslash}p{4.0cm} }
        \toprule
        \textbf{Library name} & \textbf{\#Mol} & \textbf{About} \\
        \midrule
        
        Fragment & 500,000 & It consists of a selection of fragments from the Diverse library that satisfy the rule of three (MW $<300$, LogP $<3.0$, HB donor $\le3$). \\
        \addlinespace
        
        Diverse & 1,000,000 & A standard library for Hyper Screening of 1 million compounds using HITS's own filtering. \\
        \addlinespace
        
        FDA approved & 1,100 & It consists of 1,100 FDA-approved drugs offered. \\
        \addlinespace
        
        Natural product-like & 4,200 & It consists of 4,200 molecules representing 1,000 structural motifs derived from natural products. \\
        \addlinespace
        
        Kinase focused & 65,000 & Kinase-focused library containing bioisosteric derivatives and hinge binder molecules. \\
        
        \bottomrule
    \end{tabular}
    \label{tab:hyperscreening_fixed}
\end{table}


\subsection*{Hyper Screening X}
Hyper Screening X is a service designed to discover active compounds by screening an extensive virtual library of 11 trillion molecules. Compared to conventional virtual screening methods that typically explore around 1 million compounds, Hyper Screening X offers two significant advantages: 1) increased hit discovery probability that can be dozens times higher than traditional approaches\cite{Lyu2019}, and 2) an increased likelihood of identifying novel active compounds compared to conventional commercial screening libraries. Previous research has demonstrated that virtual libraries containing billions of molecules can achieve hit discovery success rates of 20-30\%\cite{Lyu2019}. By leveraging a vast chemical space of 11 trillion molecules, Hyper Screening X provides a significantly higher probability of discovering novel, potent hit compounds compared to conventional screening methods that are limited to conventional screening libraries.

Calculating binding affinities for all 11 trillion molecules individually would require astronomical computational resources. To address this challenge, Hyper Screening X employs generative AI to identify combinations of building blocks and reaction templates capable of producing molecules with desired binding scores and properties by the user. Beyond binding affinity, the system can optimize various molecular properties including molecular weight (MW), topological polar surface area (TPSA), and LogP. Hyper Screening X utilizes state-of-the-art GFlowNet\cite{Bengio2021}-based deep learning models\cite{Seo2025} for virtual library exploration, offering the advantage of discovering more diverse molecular structures compared to reinforcement learning-based methods. Additionally, the deep learning models employed by Hyper Screening X demonstrate property optimization performance that is 5-30\% superior to similar competitive models\cite{Seo2025}.

The Hyper Screening X workflow consists of three main stages: 1) target protein and target property setting, 2) generative AI training, and 3) molecule generation upon completion of training. Training the generative AI models takes approximately 48 hours, and the subsequent generation of 100 molecules takes 30 minutes. Following molecule generation, users can request synthesis of the generated compounds through Synple Chem. After consultation with Synple Chem, the synthesized compounds can be delivered to the user. For researchers who prefer to conduct synthesis independently, Hyper Screening X also provides synthetic routes for the generated molecules. Hyper Screening is suitable when a user wants to quickly discover active compounds from their proprietary compound library or from approximately one million commercially available compounds, and then proceed to the hit-to-lead phase. If a user needs a higher hit discovery rate, simultaneous optimization of multiple properties, and identification of novel active compounds, Hyper Screening X is the appropriate choice.

\begin{figure}%[tbhp]
\centering
\includegraphics[width=1.0\linewidth]{Figures/hyperscreeningx.png}
\caption{a) Dramatic difference in library size between a conventional docking (millions of compounds) and Hyper Screening X (11 trillion compounds). b) Demonstrates the relationship between library size and hit rate. The graph shows the schematic representation that the increase of the library size to 7 trillion compounds significantly enhances the probability of identifying active compounds.}
\label{fig:hyperscreeningx}
\end{figure}

\subsection*{Hyper Design}
Hyper Design provides a feature that transforms given scaffolds to design molecules predicted to show improved biological activity against the target protein. The Hyper Design starts with predicting binding scores and binding conformations of scaffolds, then leverages this information to design molecular structures with improved binding scores. Also, Hyper Design can initiate the process from X-ray structures of small molecule-bound proteins. By registering a protein structure via its PDB ID and selecting the specific site of the ligand for modification, the system recommends new derivative structures. This approach provides inspiration for designing novel substances based on existing ligands.

Users have flexible design options, including the ability to select specific substructures of the scaffold for modification or add fragments at specific atomic positions. During molecular design, Hyper Design considers synthesizability of the generated molecules while replacing fragments or adding new ones. The fragment set used by Hyper Design was derived from the ChEMBL\cite{Zdrazil2024} compound list. In addition, the Hyper Design provides 3D structures of generated molecules, enabling users to interpret results and gain insight for further molecular design ideas.

Molecules generated through Hyper Design can be recursively subjected to further design iterations. The resulting molecules are visualized in a hierarchical flow diagram, making it easy to understand the molecular design progression. Hyper Design can serve multiple purposes, including patent circumvention of reference compounds, fragment growth in fragment-based drug design, and generation of novel molecular design ideas. You can find a case study using Hyper Design in the Case Study section.

Figure \labelcref{fig:hyperdesign} illustrates an application of HyperLab's Hyper Design capability in medicinal chemistry, specifically demonstrating hit optimization starting from a small fragment. The top-left structure shows the reference compound Imatinib, while the remaining structures represent novel derivatives generated through Hyper Design. The blue-highlighted portions in each molecule indicate the common hit fragment that served as the starting point for the design process. This scaffold acted as the initial hit molecule in the actual development process of the imatinib molecule\cite{Deininger2005}. This visualization effectively demonstrates how Hyper Design can successfully optimize an initial hit molecule into more complex molecules with structural features similar to known active compounds like Imatinib. The designs build upon the core fragment while exploring different substitution patterns and functional groups that could enhance target binding and pharmacological properties. Each variant maintains the essential hit fragment while introducing strategic modifications that could potentially improve binding affinity against the target.

\begin{figure}%[tbhp]
\centering
\includegraphics[width=1.0\linewidth]{Figures/hyperdesign.png}
\caption{Hit optimization demonstration using Hyper Design: Starting from a hit fragment (highlighted in blue), HyperLab successfully generated multiple Imatinib-like molecules by optimizing the initial hit. This capability enables efficient hit-to-lead development for drug discovery applications.}
\label{fig:hyperdesign}
\end{figure}

\subsection*{SAR analysis}

HyperLab's SAR analysis is a specialized module designed to streamline Structure-Activity Relationship (SAR) analysis, which is critical during the Hit-to-Lead and Lead Optimization stages. This feature is seamlessly integrated with "Bench", allowing users to instantly load molecular lists and begin analysis simply by specifying a Bench when creating an analysis page.

The central component of this analysis is setting a "Core" to group molecules. The system offers "Automatic Recommendation," which uses BRICS-based fragmentation to suggest core candidates based on common Murcko scaffolds, and "Manual Designation," where users can directly draw structures. Users can prioritize up to five cores for analysis. Figure \labelcref{fig:sar} illustrates the resulting R-group decomposition table. The moleculess are separated into columns by the designated Core and substituents (R1, R2, etc.) and arranged according to set alignment rules. This consistent view allows users to identify structural differences at a glance and enhances the efficiency of comparative analysis. The analysis is further deepened by the unique "Hierarchical Decomposition" feature. For example, within a complex structure, users can recursively analyze a specific sub-fragment (e.g., designating a secondary core within a substituent) to explore structural diversity in greater depth.

In addition, the system integrates HyperLab's predicted values (e.g., Hyper Binding, Hyper ADMET) with user-input experimental data. When a specific R-group column is selected, the system classifies molecules into a "Structural Series" and generates group-specific graphs. This interactive interface allows users to drag and resize molecule cards on correlation graphs to visually construct a structure-activity story. This customized analysis view can be exported as a high-resolution image for immediate use in papers or reports. 

\begin{figure}%[tbhp]
\centering
\includegraphics[width=1.0\linewidth]{Figures/sar.png}
\caption{R-group decomposition interface for SAR analysis. The molecules in the library are structurally aligned and decomposed based on the user-designated core. Each column (Core, R1, R2) maintains a consistent 2D alignment, enhancing the efficiency of visual comparative analysis.}
\label{fig:sar}
\end{figure}

\subsection*{Hyper ADME/T}
Hyper ADME/T can predict 19 key pharmacological and physicochemical properties including LogP, solubility, metabolic stability, CYP1A2 inhibition, CYP2C19 inhibition, CYP2C9 inhibition, CYP2D6 inhibition, CYP3A4 inhibition, CYP1A2 substrate, CYP2C19 substrate, CYP2C9 substrate, CYP2D6 substrate, CYP3A4 substrate, hERG inhibition, P-gp inhibition, PAMPA, druglikeness\cite{Lee2022}, synthesis complexity\cite{Kim2024}, and BBB penetration.
From the early stages of drug development, in vitro ADME/T predictions can be used to filter molecules or inform molecular design, thereby increasing the success rate of drug development. These predicted values can be easily filtered and sorted to select desired molecules. Hyper ADME/T can perform calculations for thousands of molecules and filter them based on the predicted values.

\begin{figure}%[tbhp]
\centering
\includegraphics[width=1.0\linewidth]{Figures/admet.png}
\caption{Physicochemical properties, medicinal chemistry properties, and ADME/T profile of Pirtobrutinib predicted by Hyper ADME/T.}
\label{fig:hyperadmet}
\end{figure}

\subsection*{HyperLab AI assistant}
HyperLab features an AI assistant that enhances the efficiency of drug discovery researchers. Unlike conventional LLMs, HyperLab's AI assistant is a specialized AI agent independently developed by HITS to perform drug discovery-specific workflows. HyperLab's AI assistant is powered by Claude 3.7 provided from AWS. It offers capabilities such as searching specialized drug discovery databases like PubChem\cite{Kim2025}, ChEMBL\cite{Zdrazil2024}, and RCSB PDB\cite{Zardecki2022}, conducting web searches, coding with molecule-specific libraries, retrieving clinical trial information, visualizing graphs, and providing guidance on HyperLab usage.
For example, after uploading data to HyperLab, a user can simply request "Draw a correlation between Hyper Binding score and experimental values," and the system will generate the corresponding graph. The HyperLab AI assistant is accessible from anywhere within HyperLab and provides higher quality responses and consistency compared to general-purpose chat AIs like ChatGPT when it comes to drug discovery tasks.

Figure \labelcref{fig:AIassistant} shows an example response of HyperLab AI Assistant providing a comprehensive analysis of BI-2852, a KRAS inhibitor developed by Boehringer Ingelheim. The AI assistant explains that as of April 2025, BI-2852 has not entered clinical trials and remains primarily a research tool compound available through Boehringer Ingelheim's opnMe platform. The response details BI-2852's mechanism of action, including its binding to KRAS G12D, inhibition properties, and preclinical activity profile.

\begin{figure}%[tbhp]
\centering
\includegraphics[width=1.0\linewidth]{Figures/BI2852}
\caption{An example of how the HyperLab AI assistant would respond when a user asks about the clinical stage of BI-2852.}
\label{fig:AIassistant}
\end{figure}


\section*{Benchmark results}
\subsection*{Pose prediction accuracy of protein-ligand complex}
To assess the pose prediction performance of various docking and structure-based modeling approaches, we conducted experiments on the PoseBuster v2\cite{Buttenschoen2024} (PB-valid) benchmark. This benchmark is a curated dataset specifically designed to evaluate the accuracy of ligand binding pose predictions across a wide range of protein targets and chemical scaffolds, making it a rigorous testbed for assessing generalizability and robustness.

As shown in Figure \labelcref{fig:benchmark1}, rigid docking methods such as Diffdock\cite{Corso2023} and Vina\cite{Eberhardt2021} demonstrated limited accuracy (13\% and 58\%, respectively), likely due to their reliance on fixed receptor conformations and simplistic scoring functions. Chai Discovery\cite{Boitreaud2024} improved upon these methods (66\%), while Hyper binding co-folding showed notable gains with 68\% accuracy. When provided with binding site information, our method reached 77\% accuracy, closely approaching the performance of AlphaFold3\cite{Abramson2024} (84\%) and Boltz2\cite{Passaro2025} (78\%).

\begin{figure}%[tbhp]
\centering
\includegraphics[width=1.0\linewidth]{Figures/benchmark1.png}
\caption{Binding pose prediction accuracy on the PoseBuster v2 (PB-valid) benchmark. Hyper binding co-folding (with binding site) achieves 77\% accuracy, outperforming traditional docking methods and approaching the accuracy of AlphaFold3 (84\%).}
\label{fig:benchmark1}
\end{figure}

Note that unlike Vina and other docking approaches, Hyper binding co-folding does not utilize any 3D structural information of the target protein during inference. Despite this, it achieves comparable or better accuracy, highlighting the power of end-to-end complex co-folding in learning spatial constraints directly from sequence and ligand input. Importantly, our Hyper binding co-folding method offers a significant advantage in inference speed. Importantly, while AlphaFold3 requires approximately 15 minutes\cite{Voitsitskyi2024} per complex with a RTX 3060 GPU, our approach completes the co-folding calculation for the same complex in just 3 minutes by using vast cloud resources. This five-fold speed advantage makes Hyper Binding co-folding a highly practical and scalable solution for virtual screening campaigns and rapid design-test-learn cycles, where computational throughput is a critical bottleneck.

\subsection*{Binding affinity prediction accuracy of protein-ligand complex}

Predicting the activity of derivatives sharing a common scaffold is an extremely valuable technique in hit-to-lead and lead optimization processes. However, this is a particularly challenging task because the activity differences between derivatives are often very subtle, requiring precise predictions, and small structural changes can sometimes have significant impacts on activity values. We performed benchmarking on this task using two benchmark datasets (referred to as FEP benchmark1\cite{Wang2015} and FEP benchmark2\cite{Schindler2020}).

Figure \labelcref{fig:benchmark2} illustrates the Pearson correlation coefficient achieved by five different models on two distinct datasets, FEP dataset1 and FEP dataset2. The models evaluated are Hyper Binding, Luminet\cite{Su2025}, Genscore\cite{Shen2023}, Glide SP\cite{Friesner2004}, and Vina.
On FEP dataset1, Hyper Binding demonstrated the highest performance with a Pearson correlation coefficient of 0.70. Luminet followed with a coefficient of 0.65, and Genscore achieved 0.57. Glide SP and Vina showed lower performance on this dataset, with coefficients of 0.29 and 0.25, respectively.

For FEP dataset2, Hyper Binding again achieved the highest Pearson correlation coefficient at 0.53, followed closely by Genscore at 0.52. Luminet's performance was 0.46. Similar to FEP dataset1, docking methods such as Glide SP and Vina had the lowest scores on FEP dataset2, with coefficients of 0.24 and 0.34, respectively. Overall, Hyper Binding consistently outperformed the both deep learning models and physics based docking methods on both datasets. 

\begin{figure}%[tbhp]
\centering
\includegraphics[width=1.0\linewidth]{Figures/benchmark2.png}
\caption{The bar chart compares the Pearson correlation coefficients of Hyper Binding against other deep learning models (Luminet, Genscore) and traditional physics-based docking methods (Glide SP, Vina). On both FEP dataset1 and FEP dataset2, Hyper Binding consistently achieved the highest correlation, demonstrating its superior accuracy in predicting binding affinity.}
\label{fig:benchmark2}
\end{figure}

\section*{Case study: AI-enabled screening and design with HyperLab}
HyperLab was created to compress the early phases of drug discovery by combining ultra-fast virtual screening (“Hyper Screening”) with an AI-guided compound-design module (“Hyper Design”). To evaluate the platform’s technical performance, we conducted an internal validation study using a pipeline developed in-house.
In the Hyper Screening stage, we selected the PDB structure of the target protein and performed Hyper Binding calculations against a diverse compound library. Based on binding scores, we directly selected 52 top-ranked compounds for in vitro testing. The entire virtual screening process was completed within 24 hours. In vitro assay results (Figure \labelcref{fig:casestudy_1}) identified five compounds showing >40\% inhibition, corresponding to a hit rate of approximately 9\%. The IC$_{50}$ values of the identified compounds, including the reference compound H00112 (IC$_{50}$=1319 nM), ranged from 70 to 600 nM.

\begin{figure}%[tbhp]
\centering
\includegraphics[width=1.0\linewidth]{Figures/casestudy_1.jpg}
\caption{Hit validation of top-ranked compounds from Hyper Screening. Dose–response curves for the six hit compounds identified in, measured in a kinase inhibition assay. Curves were fitted using a standard sigmoid model to determine IC$_{50}$ values. All data are expressed as mean ± SD from duplicate experiments, relative to the DMSO control. Assays were performed by Eurofins Discovery using the KinaseProfiler LeadHunter radiometric platform.}
\label{fig:casestudy_1}
\end{figure}

Using the Hyper Design, we conducted a part-by-part replacement of the reference compound to generate structurally novel derivatives. Compound selection was guided by multiple criteria, including improvement in predicted binding scores, retention of key molecular interactions with the target protein, and structural novelty with a particular emphasis on avoiding patent-protected motifs of the reference compound. Among the 14 candidate compounds selected through this process, 10 (71\%) were assessed to be synthetically accessible based on human inspection. Five of these were synthesized early and subsequently subjected to in vitro assays. The three compounds exhibited over 75\% inhibition at a concentration of 1 $\mu$M, with IC$_{50}$ values ranging from 200 to 400 nM. These results are shown in Figure \cref{fig:casestudy_2}. 

H00505 emerged as the most potent compound among those tested, exhibiting inhibitory activity comparable to that of the reference compound H00369. This finding prompted further mechanistic studies to evaluate its engagement with the relevant signaling pathway. In the pathway-specific assay, measured by the ARE luciferase reporter system (Figure \cref{fig:casestudy_3}), H00505 demonstrated activity equal to or greater than that of the reference compound, highlighting its potential as a promising lead candidate for further optimization.

\begin{figure}%[tbhp]
\centering
\includegraphics[width=1.0\linewidth]{Figures/casestudy_2.png}
\caption{Inhibitory activity of synthesized compounds against the target protein. a) Single-dose assay at 1 μM showing the percentage activity of the target protein relative to the DMSO control (set as 100\%). H00369 is the reference compound, and all others are newly synthesized derivatives. b) Dose–response curves of selected compounds (H00502, H00505, H00506, and H00369) plotted as percentage activity versus log$_{10}$[compound] ($\mu$M). All data are expressed as mean ± SD from duplicate experiments, relative to the DMSO control. Assays were performed by BPS Bioscience}
\label{fig:casestudy_2}
\end{figure}

\begin{figure}%[tbhp]
\centering
\includegraphics[width=0.4\linewidth]{Figures/casestudy_3.jpg}
\caption{The ARE transcriptional activity of H00505. ARE–luciferase–HepG2 cells were treated for 18 h with 10$\mu$M of H00505 and the reference compound, and luciferase activity was measured using the ONE-Step™ Luciferase Assay System (BPS Bioscience). H00505 induced a significantly greater fold increase in ARE activity compared to both the DMSO control and the reference compound, indicating enhanced activation of the antioxidant response pathway. Data are presented as mean ± SD from triplicate experiments, relative to the DMSO control. p values were determined using an unpaired two-tailed t-test. p<0.05 (*), p<0.001 (***).}
\label{fig:casestudy_3}
\end{figure}

Taken together, these results demonstrate that HyperLab efficiently identifies novel hit compounds and converts in-silico designs into experimentally validated leads with nanomolar potency, supporting its application as a scalable and cost-effective platform for target-based drug discovery.

Conventional high-throughput screening (HTS) campaigns typically interrogate $\geq$105 compounds in single-concentration assays, with subsequent confirmatory titrations to obtain IC$_{50}$ values, and report primary hit rates in the low single-digit percent range. In contrast, Hyper Screening identified 6 actives (11.5\%) from only 52 candidates, while Hyper Design yielded 3 actives out of 5 synthesized (60\%), thereby introducing structural differentiation without loss of potency. Assuming standard assay workflows (duplicate single-point and 10-point duplicate titrations), this corresponds to a reduction from ~200,000 wells in a representative HTS to ~100 wells in our pipelines. These results demonstrate the efficiency gains achievable by computational triage and AI-guided motif design in reducing both time and cost while maintaining pharmacological relevance.

\section*{Conclusion}
HyperLab represents a significant advancement in AI-driven drug discovery platforms, addressing the critical need for accessible computational tools for drug discovery researchers. By combining state-of-the-art AI models with an intuitive user interface, HyperLab democratizes access to sophisticated drug discovery capabilities that were previously available only to computational specialists. The platform's comprehensive suite of tools—Hyper Binding, Hyper Design, Hyper Screening, and Hyper ADME/T—covers the entire spectrum of structure-based drug discovery needs, from initial hit identification to lead optimization.

Our benchmark results demonstrate HyperLab's superior performance compared to conventional methods, with Hyper Binding achieving 77\% accuracy in binding pose prediction and consistently outperforming both deep learning models and physics-based docking methods in binding affinity prediction across multiple datasets. These performance metrics, combined with the platform's user-friendly interface and specialized AI assistant, position HyperLab as a transformative tool for accelerating drug discovery processes.

The practical power of this integrated platform was validated through an internal case study, which progressed from initial high-throughput screening to the experimental confirmation of a novel lead compound with nanomolar potency (IC$_{50}$ values of 200-400 nM). 
This achievement demonstrates HyperLab's real-world capability to dramatically compress discovery timelines and efficiently convert in-silico designs into experimentally validated leads. As variety of AI methods continue to play an increasingly vital role in pharmaceutical research, HyperLab stands poised to significantly reduce the time, cost, and expertise barriers associated with modern drug discovery efforts.


%% \begin{acknowledgements}
%%  \blindtext
%% \end{acknowledgements}
\section*{Revision History}

\begin{description} % 옵션 제거

    \item[2026-01] 
    SAR feature release update. \\

    \item[2025-09] 
    Initial release of the HyperLab platform documentation. \\

\end{description}

\vspace{1em} % 표와 다음 본문 사이의 간격 추가

\section*{Bibliography}
\bibliography{zHenriquesLab-Mendeley}

%% You can use these special %TC: tags to ignore certain parts of the text.
%TC:ignore
%the command above ignores this section for word count


\end{document}
