## Supplementary material for "Accelerating Drug Discovery with HyperLab: An Easy-to-Use AI-Driven Platform": all latex file: zHenriquesLab-Mendeley.bib

@article{Seo2025,
   abstract = {Generative models in drug discovery have recently gained attention as efficient alternatives to brute-force virtual screening. However, most existing models do not account for synthesizability, limiting their practical use in real-world scenarios. In this paper, we propose RxnFlow, which sequentially assembles molecules using predefined molecular building blocks and chemical reaction templates to constrain the synthetic chemical pathway. We then train on this sequential generating process with the objective of generative flow networks (GFlowNets) to generate both highly rewarded and diverse molecules. To mitigate the large action space of synthetic pathways in GFlowNets, we implement a novel action space subsampling method. This enables RxnFlow to learn generative flows over extensive action spaces comprising combinations of 1.2 million building blocks and 71 reaction templates without significant computational overhead. Additionally, RxnFlow can employ modified or expanded action spaces for generation without retraining, allowing for the introduction of additional objectives or the incorporation of newly discovered building blocks. We experimentally demonstrate that RxnFlow outperforms existing reaction-based and fragment-based models in pocket-specific optimization across various target pockets. Furthermore, RxnFlow achieves state-of-the-art performance on CrossDocked2020 for pocket-conditional generation, with an average Vina score of -8.85 kcal/mol and 34.8% synthesizability.},
   author = {Seonghwan Seo and Minsu Kim and Tony Shen and Martin Ester and Jinkyoo Park and Sungsoo Ahn and Woo Youn Kim},
   month = {3},
   title = {Generative Flows on Synthetic Pathway for Drug Design},
   year = {2025}
}
@article{Su2025,
   abstract = {Robust protein–ligand interaction modeling through integrating physical laws and geometric knowledge for absolute binding free energy calculation.},
   author = {Qun Su and Jike Wang and Qiaolin Gou and Renling Hu and Linlong Jiang and Hui Zhang and Tianyue Wang and Yifei Liu and Chao Shen and Yu Kang and Chang-Yu Hsieh and Tingjun Hou},
   doi = {10.1039/D4SC07405J},
   issn = {2041-6520},
   issue = {12},
   journal = {Chemical Science},
   pages = {5043-5057},
   title = {Robust protein–ligand interaction modeling through integrating physical laws and geometric knowledge for absolute binding free energy calculation},
   volume = {16},
   year = {2025}
}
@article{Abramson2024,
   abstract = { The introduction of AlphaFold 2 1 has spurred a revolution in modelling the structure of proteins and their interactions, enabling a huge range of applications in protein modelling and design 2–6 . Here we describe our AlphaFold 3 model with a substantially updated diffusion-based architecture that is capable of predicting the joint structure of complexes including proteins, nucleic acids, small molecules, ions and modified residues. The new AlphaFold model demonstrates substantially improved accuracy over many previous specialized tools: far greater accuracy for protein–ligand interactions compared with state-of-the-art docking tools, much higher accuracy for protein–nucleic acid interactions compared with nucleic-acid-specific predictors and substantially higher antibody–antigen prediction accuracy compared with AlphaFold-Multimer v.2.3 7,8 . Together, these results show that high-accuracy modelling across biomolecular space is possible within a single unified deep-learning framework. },
   author = {Josh Abramson and Jonas Adler and Jack Dunger and Richard Evans and Tim Green and Alexander Pritzel and Olaf Ronneberger and Lindsay Willmore and Andrew J. Ballard and Joshua Bambrick and Sebastian W. Bodenstein and David A. Evans and Chia-Chun Hung and Michael O’Neill and David Reiman and Kathryn Tunyasuvunakool and Zachary Wu and Akvilė Žemgulytė and Eirini Arvaniti and Charles Beattie and Ottavia Bertolli and Alex Bridgland and Alexey Cherepanov and Miles Congreve and Alexander I. Cowen-Rivers and Andrew Cowie and Michael Figurnov and Fabian B. Fuchs and Hannah Gladman and Rishub Jain and Yousuf A. Khan and Caroline M. R. Low and Kuba Perlin and Anna Potapenko and Pascal Savy and Sukhdeep Singh and Adrian Stecula and Ashok Thillaisundaram and Catherine Tong and Sergei Yakneen and Ellen D. Zhong and Michal Zielinski and Augustin Žídek and Victor Bapst and Pushmeet Kohli and Max Jaderberg and Demis Hassabis and John M. Jumper},
   doi = {10.1038/s41586-024-07487-w},
   issn = {0028-0836},
   issue = {8016},
   journal = {Nature},
   month = {6},
   pages = {493-500},
   pmid = {38718835},
   publisher = {Nature Research},
   title = {Accurate structure prediction of biomolecular interactions with AlphaFold 3},
   volume = {630},
   url = {https://www.nature.com/articles/s41586-024-07487-w},
   year = {2024}
}
@article{Moon2024,
   abstract = {PIGNet2, a versatile protein–ligand interaction prediction model that performs well in both molecule identification and optimization, demonstrates its potential in early-stage drug discovery.},
   author = {Seokhyun Moon and Sang-Yeon Hwang and Jaechang Lim and Woo Youn Kim},
   doi = {10.1039/D3DD00149K},
   issn = {2635-098X},
   issue = {2},
   journal = {Digital Discovery},
   pages = {287-299},
   title = {PIGNet2: a versatile deep learning-based protein–ligand interaction prediction model for binding affinity scoring and virtual screening},
   volume = {3},
   year = {2024}
}
@article{Buttenschoen2024,
   abstract = {PoseBusters assesses molecular poses using steric and energetic criteria. We find that classical protein-ligand docking tools currently still outperform deep learning-based methods.},
   author = {Martin Buttenschoen and Garrett M. Morris and Charlotte M. Deane},
   doi = {10.1039/D3SC04185A},
   issn = {2041-6520},
   issue = {9},
   journal = {Chemical Science},
   pages = {3130-3139},
   title = {PoseBusters: AI-based docking methods fail to generate physically valid poses or generalise to novel sequences},
   volume = {15},
   year = {2024}
}
@misc{Boitreaud2024,
   abstract = {We introduce Chai-1, a multi-modal foundation model for molecular structure prediction that performs at the state-of-the-art across a variety of tasks relevant to drug discovery. Chai-1 can optionally be prompted with experimental restraints (e.g. derived from wet-lab data) which boosts performance by double-digit percentage points. Chai-1 can also be run in single-sequence mode with-out MSAs while preserving most of its performance. We release Chai-1 model weights and inference code as a Python package for non-commercial use and via a web interface where it can be used for free including for commercial drug discovery purposes.},
   author = {Jacques Boitreaud and Jack Dent and Matthew McPartlon and Joshua Meier and Vinicius Reis and Alex Rogozhnikov and Kevin Wu},
   doi = {10.1101/2024.10.10.615955},
   month = {10},
   title = {Chai-1: Decoding the molecular interactions of life},
   year = {2024}
}
@article{Kim2024,
   author = {Hyeongwoo Kim and Kyunghoon Lee and Chansu Kim and Jaechang Lim and Woo Youn Kim},
   doi = {10.1021/acs.jcim.3c01134},
   issn = {1549-9596},
   issue = {7},
   journal = {Journal of Chemical Information and Modeling},
   month = {4},
   pages = {2432-2444},
   title = {DFRscore: Deep Learning-Based Scoring of Synthetic Complexity with Drug-Focused Retrosynthetic Analysis for High-Throughput Virtual Screening},
   volume = {64},
   year = {2024}
}
@article{Corso2023,
   abstract = {Predicting the binding structure of a small molecule ligand to a protein -- a task known as molecular docking -- is critical to drug design. Recent deep learning methods that treat docking as a regression problem have decreased runtime compared to traditional search-based methods but have yet to offer substantial improvements in accuracy. We instead frame molecular docking as a generative modeling problem and develop DiffDock, a diffusion generative model over the non-Euclidean manifold of ligand poses. To do so, we map this manifold to the product space of the degrees of freedom (translational, rotational, and torsional) involved in docking and develop an efficient diffusion process on this space. Empirically, DiffDock obtains a 38% top-1 success rate (RMSD<2A) on PDBBind, significantly outperforming the previous state-of-the-art of traditional docking (23%) and deep learning (20%) methods. Moreover, while previous methods are not able to dock on computationally folded structures (maximum accuracy 10.4%), DiffDock maintains significantly higher precision (21.7%). Finally, DiffDock has fast inference times and provides confidence estimates with high selective accuracy.},
   author = {Gabriele Corso and Hannes Stärk and Bowen Jing and Regina Barzilay and Tommi Jaakkola},
   month = {2},
   title = {DiffDock: Diffusion Steps, Twists, and Turns for Molecular Docking},
   year = {2023}
}
@article{Shen2023,
   abstract = {A generalized protein–ligand scoring framework with balanced scoring, docking, ranking and screening powers are developed here.},
   author = {Chao Shen and Xujun Zhang and Chang-Yu Hsieh and Yafeng Deng and Dong Wang and Lei Xu and Jian Wu and Dan Li and Yu Kang and Tingjun Hou and Peichen Pan},
   doi = {10.1039/D3SC02044D},
   issn = {2041-6520},
   issue = {30},
   journal = {Chemical Science},
   pages = {8129-8146},
   title = {A generalized protein–ligand scoring framework with balanced scoring, docking, ranking and screening powers},
   volume = {14},
   year = {2023}
}
@article{Lee2022,
   abstract = {A new quantification method of drug-likeness based on unsupervised learning. The method only uses drug molecules as training set without any non-drug-like molecules.},
   author = {Kyunghoon Lee and Jinho Jang and Seonghwan Seo and Jaechang Lim and Woo Youn Kim},
   doi = {10.1039/D1SC05248A},
   issn = {2041-6520},
   issue = {2},
   journal = {Chemical Science},
   pages = {554-565},
   title = {Drug-likeness scoring based on unsupervised learning},
   volume = {13},
   year = {2022}
}
@article{Bengio2021,
   abstract = {This paper is about the problem of learning a stochastic policy for generating an object (like a molecular graph) from a sequence of actions, such that the probability of generating an object is proportional to a given positive reward for that object. Whereas standard return maximization tends to converge to a single return-maximizing sequence, there are cases where we would like to sample a diverse set of high-return solutions. These arise, for example, in black-box function optimization when few rounds are possible, each with large batches of queries, where the batches should be diverse, e.g., in the design of new molecules. One can also see this as a problem of approximately converting an energy function to a generative distribution. While MCMC methods can achieve that, they are expensive and generally only perform local exploration. Instead, training a generative policy amortizes the cost of search during training and yields to fast generation. Using insights from Temporal Difference learning, we propose GFlowNet, based on a view of the generative process as a flow network, making it possible to handle the tricky case where different trajectories can yield the same final state, e.g., there are many ways to sequentially add atoms to generate some molecular graph. We cast the set of trajectories as a flow and convert the flow consistency equations into a learning objective, akin to the casting of the Bellman equations into Temporal Difference methods. We prove that any global minimum of the proposed objectives yields a policy which samples from the desired distribution, and demonstrate the improved performance and diversity of GFlowNet on a simple domain where there are many modes to the reward function, and on a molecule synthesis task.},
   author = {Emmanuel Bengio and Moksh Jain and Maksym Korablyov and Doina Precup and Yoshua Bengio},
   month = {11},
   title = {Flow Network based Generative Models for Non-Iterative Diverse Candidate Generation},
   year = {2021}
}
@article{Eberhardt2021,
   author = {Jerome Eberhardt and Diogo Santos-Martins and Andreas F. Tillack and Stefano Forli},
   doi = {10.1021/acs.jcim.1c00203},
   issn = {1549-9596},
   issue = {8},
   journal = {Journal of Chemical Information and Modeling},
   month = {8},
   pages = {3891-3898},
   title = {AutoDock Vina 1.2.0: New Docking Methods, Expanded Force Field, and Python Bindings},
   volume = {61},
   year = {2021}
}
@article{Schindler2020,
   author = {Christina E. M. Schindler and Hannah Baumann and Andreas Blum and Dietrich Böse and Hans-Peter Buchstaller and Lars Burgdorf and Daniel Cappel and Eugene Chekler and Paul Czodrowski and Dieter Dorsch and Merveille K. I. Eguida and Bruce Follows and Thomas Fuchß and Ulrich Grädler and Jakub Gunera and Theresa Johnson and Catherine Jorand Lebrun and Srinivasa Karra and Markus Klein and Tim Knehans and Lisa Koetzner and Mireille Krier and Matthias Leiendecker and Birgitta Leuthner and Liwei Li and Igor Mochalkin and Djordje Musil and Constantin Neagu and Friedrich Rippmann and Kai Schiemann and Robert Schulz and Thomas Steinbrecher and Eva-Maria Tanzer and Andrea Unzue Lopez and Ariele Viacava Follis and Ansgar Wegener and Daniel Kuhn},
   doi = {10.1021/acs.jcim.0c00900},
   issn = {1549-9596},
   issue = {11},
   journal = {Journal of Chemical Information and Modeling},
   month = {11},
   pages = {5457-5474},
   title = {Large-Scale Assessment of Binding Free Energy Calculations in Active Drug Discovery Projects},
   volume = {60},
   year = {2020}
}
@article{Wang2015,
   abstract = {Designing tight-binding ligands is a primary objective of small-molecule drug discovery. Over the past few decades, free-energy calculations have benefited from improved force fields and sampling algorithms, as well as the advent of low-cost parallel computing. However, it has proven to be challenging to reliably achieve the level of accuracy that would be needed to guide lead optimization (5× in binding affinity) for a wide range of ligands and protein targets. Not surprisingly, widespread commercial application of free-energy simulations has been limited due to the lack of large-scale validation coupled with the technical challenges traditionally associated with running these types of calculations. Here, we report an approach that achieves an unprecedented level of accuracy across a broad range of target classes and ligands, with retrospective results encompassing 200 ligands and a wide variety of chemical perturbations, many of which involve significant changes in ligand chemical structures. In addition, we have applied the method in prospective drug discovery projects and found a significant improvement in the quality of the compounds synthesized that have been predicted to be potent. Compounds predicted to be potent by this approach have a substantial reduction in false positives relative to compounds synthesized on the basis of other computational or medicinal chemistry approaches. Furthermore, the results are consistent with those obtained from our retrospective studies, demonstrating the robustness and broad range of applicability of this approach, which can be used to drive decisions in lead optimization.},
   author = {Lingle Wang and Yujie Wu and Yuqing Deng and Byungchan Kim and Levi Pierce and Goran Krilov and Dmitry Lupyan and Shaughnessy Robinson and Markus K. Dahlgren and Jeremy Greenwood and Donna L. Romero and Craig Masse and Jennifer L. Knight and Thomas Steinbrecher and Thijs Beuming and Wolfgang Damm and Ed Harder and Woody Sherman and Mark Brewer and Ron Wester and Mark Murcko and Leah Frye and Ramy Farid and Teng Lin and David L. Mobley and William L. Jorgensen and Bruce J. Berne and Richard A. Friesner and Robert Abel},
   doi = {10.1021/ja512751q},
   issn = {0002-7863},
   issue = {7},
   journal = {Journal of the American Chemical Society},
   month = {2},
   pages = {2695-2703},
   publisher = {American Chemical Society},
   title = {Accurate and Reliable Prediction of Relative Ligand Binding Potency in Prospective Drug Discovery by Way of a Modern Free-Energy Calculation Protocol and Force Field},
   volume = {137},
   url = {https://pubs.acs.org/doi/10.1021/ja512751q},
   year = {2015}
}
@article{Friesner2004,
   author = {Richard A. Friesner and Jay L. Banks and Robert B. Murphy and Thomas A. Halgren and Jasna J. Klicic and Daniel T. Mainz and Matthew P. Repasky and Eric H. Knoll and Mee Shelley and Jason K. Perry and David E. Shaw and Perry Francis and Peter S. Shenkin},
   doi = {10.1021/jm0306430},
   issn = {0022-2623},
   issue = {7},
   journal = {Journal of Medicinal Chemistry},
   month = {3},
   pages = {1739-1749},
   title = {Glide:  A New Approach for Rapid, Accurate Docking and Scoring. 1. Method and Assessment of Docking Accuracy},
   volume = {47},
   year = {2004}
}
@article{Varadi2024,
   abstract = {The AlphaFold Database Protein Structure Database (AlphaFold DB, https://alphafold.ebi.ac.uk) has significantly impacted structural biology by amassing over 214 million predicted protein structures, expanding from the initial 300k structures released in 2021. Enabled by the groundbreaking AlphaFold2 artificial intelligence (AI) system, the predictions archived in AlphaFold DB have been integrated into primary data resources such as PDB, UniProt, Ensembl, InterPro and MobiDB. Our manuscript details subsequent enhancements in data archiving, covering successive releases encompassing model organisms, global health proteomes, Swiss-Prot integration, and a host of curated protein datasets. We detail the data access mechanisms of AlphaFold DB, from direct file access via FTP to advanced queries using Google Cloud Public Datasets and the programmatic access endpoints of the database. We also discuss the improvements and services added since its initial release, including enhancements to the Predicted Aligned Error viewer, customisation options for the 3D viewer, and improvements in the search engine of AlphaFold DB.},
   author = {Mihaly Varadi and Damian Bertoni and Paulyna Magana and Urmila Paramval and Ivanna Pidruchna and Malarvizhi Radhakrishnan and Maxim Tsenkov and Sreenath Nair and Milot Mirdita and Jingi Yeo and Oleg Kovalevskiy and Kathryn Tunyasuvunakool and Agata Laydon and Augustin Žídek and Hamish Tomlinson and Dhavanthi Hariharan and Josh Abrahamson and Tim Green and John Jumper and Ewan Birney and Martin Steinegger and Demis Hassabis and Sameer Velankar},
   doi = {10.1093/nar/gkad1011},
   issn = {0305-1048},
   issue = {D1},
   journal = {Nucleic Acids Research},
   month = {1},
   pages = {D368-D375},
   title = {AlphaFold Protein Structure Database in 2024: providing structure coverage for over 214 million protein sequences},
   volume = {52},
   year = {2024}
}
@article{Buttenschoen2024,
   abstract = {PoseBusters assesses molecular poses using steric and energetic criteria. We find that classical protein-ligand docking tools currently still outperform deep learning-based methods.},
   author = {Martin Buttenschoen and Garrett M. Morris and Charlotte M. Deane},
   doi = {10.1039/D3SC04185A},
   issn = {2041-6520},
   issue = {9},
   journal = {Chemical Science},
   pages = {3130-3139},
   title = {PoseBusters: AI-based docking methods fail to generate physically valid poses or generalise to novel sequences},
   volume = {15},
   year = {2024}
}
@article{Zdrazil2024,
   abstract = {ChEMBL (https://www.ebi.ac.uk/chembl/) is a manually curated, high-quality, large-scale, open, FAIR and Global Core Biodata Resource of bioactive molecules with drug-like properties, previously described in the 2012, 2014, 2017 and 2019 Nucleic Acids Research Database Issues. Since its introduction in 2009, ChEMBL’s content has changed dramatically in size and diversity of data types. Through incorporation of multiple new datasets from depositors since the 2019 update, ChEMBL now contains slightly more bioactivity data from deposited data vs data extracted from literature. In collaboration with the EUbOPEN consortium, chemical probe data is now regularly deposited into ChEMBL. Release 27 made curated data available for compounds screened for potential anti-SARS-CoV-2 activity from several large-scale drug repurposing screens. In addition, new patent bioactivity data have been added to the latest ChEMBL releases, and various new features have been incorporated, including a Natural Product likeness score, updated flags for Natural Products, a new flag for Chemical Probes, and the initial annotation of the action type for ∼270 000 bioactivity measurements.},
   author = {Barbara Zdrazil and Eloy Felix and Fiona Hunter and Emma J Manners and James Blackshaw and Sybilla Corbett and Marleen de Veij and Harris Ioannidis and David Mendez Lopez and Juan F Mosquera and Maria Paula Magarinos and Nicolas Bosc and Ricardo Arcila and Tevfik Kizilören and Anna Gaulton and A Patrícia Bento and Melissa F Adasme and Peter Monecke and Gregory A Landrum and Andrew R Leach},
   doi = {10.1093/nar/gkad1004},
   issn = {0305-1048},
   issue = {D1},
   journal = {Nucleic Acids Research},
   month = {1},
   pages = {D1180-D1192},
   title = {The ChEMBL Database in 2023: a drug discovery platform spanning multiple bioactivity data types and time periods},
   volume = {52},
   year = {2024}
}
@article{Deininger2005,
   abstract = {Imatinib has revolutionized drug therapy of chronic myeloid leukemia (CML). Preclinical studies were promising but the results of clinical trials by far exceeded expectations. Responses in chronic phase are unprecedented, with rates of complete cytogenetic response (CCR) of more than 40% in patients after failure of interferon-α (IFN) and more than 80% in newly diagnosed patients, a level of efficacy that led to regulatory approval in record time. While most of these responses are stable, resistance to treatment after an initial response is common in more advanced phases of the disease. Mutations in the kinase domain (KD) of BCR-ABL that impair imatinib binding have been identified as the leading cause of resistance. Patients with CCR who achieve a profound reduction of BCR-ABL mRNA have a very low risk of disease progression. However, residual disease usually remains detectable with reverse transcription–polymerase chain reaction (RT-PCR), indicating that disease eradication may pose a significant challenge. The mechanisms underlying the persistence of minimal residual disease are unknown. In this manuscript, we review the preclinical and clinical development of imatinib for the therapy of CML, resistance and strategies that may help to eliminate resistant or residual leukemia.},
   author = {Michael Deininger and Elisabeth Buchdunger and Brian J. Druker},
   doi = {10.1182/blood-2004-08-3097},
   issn = {0006-4971},
   issue = {7},
   journal = {Blood},
   month = {4},
   pages = {2640-2653},
   title = {The development of imatinib as a therapeutic agent for chronic myeloid leukemia},
   volume = {105},
   year = {2005}
}
@article{Lyu2019,
   author = {Jiankun Lyu and Sheng Wang and Trent E. Balius and Isha Singh and Anat Levit and Yurii S. Moroz and Matthew J. O’Meara and Tao Che and Enkhjargal Algaa and Kateryna Tolmachova and Andrey A. Tolmachev and Brian K. Shoichet and Bryan L. Roth and John J. Irwin},
   doi = {10.1038/s41586-019-0917-9},
   issn = {0028-0836},
   issue = {7743},
   journal = {Nature},
   month = {2},
   pages = {224-229},
   title = {Ultra-large library docking for discovering new chemotypes},
   volume = {566},
   year = {2019}
}
@article{Kim2025,
   abstract = {PubChem (https://pubchem.ncbi.nlm.nih.gov) is a large and highly-integrated public chemical database resource at NIH. In the past two years, significant updates were made to PubChem. With additions from over 130 new sources, PubChem contains &gt;1000 data sources, 119 million compounds, 322 million substances and 295 million bioactivities. New interfaces, such as the consolidated literature panel and the patent knowledge panel, were developed. The consolidated literature panel combines all references about a compound into a single list, allowing users to easily find, sort, and export all relevant articles for a chemical in one place. The patent knowledge panels for a given query chemical or gene display chemicals, genes, and diseases co-mentioned with the query in patent documents, helping users to explore relationships between co-occurring entities within patent documents. PubChemRDF was expanded to include the co-occurrence data underlying the literature knowledge panel, enabling users to exploit semantic web technologies to explore entity relationships based on the co-occurrences in the scientific literature. The usability and accessibility of information on chemicals with non-discrete structures (e.g. biologics, minerals, polymers, UVCBs and glycans) were greatly improved with dedicated web pages that provide a comprehensive view of all available information in PubChem for these chemicals.},
   author = {Sunghwan Kim and Jie Chen and Tiejun Cheng and Asta Gindulyte and Jia He and Siqian He and Qingliang Li and Benjamin A Shoemaker and Paul A Thiessen and Bo Yu and Leonid Zaslavsky and Jian Zhang and Evan E Bolton},
   doi = {10.1093/nar/gkae1059},
   issn = {0305-1048},
   issue = {D1},
   journal = {Nucleic Acids Research},
   month = {1},
   pages = {D1516-D1525},
   title = {PubChem 2025 update},
   volume = {53},
   year = {2025}
}
@article{Zardecki2022,
   abstract = { The Protein Data Bank (PDB) archive is a rich source of information in the form of atomic‐level three‐dimensional (3D) structures of biomolecules experimentally determined using macromolecular crystallography, nuclear magnetic resonance (NMR) spectroscopy, and electron microscopy (3DEM). Originally established in 1971 as a resource for protein crystallographers to freely exchange data, today PDB data drive research and education across scientific disciplines. In 2011, the online portal PDB‐101 was launched to support teachers, students, and the general public in PDB archive exploration ( pdb101.rcsb.org ). Maintained by the Research Collaboratory for Structural Bioinformatics PDB, PDB‐101 aims to help train the next generation of PDB users and to promote the overall importance of structural biology and protein science to nonexperts. Regularly published features include the highly popular Molecule of the Month series, 3D model activities, molecular animation videos, and educational curricula. Materials are organized into various categories (Health and Disease, Molecules of Life, Biotech and Nanotech, and Structures and Structure Determination) and searchable by keyword. A biennial health focus frames new resource creation and provides topics for annual video challenges for high school students. Web analytics document that PDB‐101 materials relating to fundamental topics (e.g., hemoglobin, catalase) are highly accessed year‐on‐year. In addition, PDB‐101 materials created in response to topical health matters (e.g., Zika, measles, coronavirus) are well received. PDB‐101 shows how learning about the diverse shapes and functions of PDB structures promotes understanding of all aspects of biology, from the central dogma of biology to health and disease to biological energy. },
   author = {Christine Zardecki and Shuchismita Dutta and David S. Goodsell and Robert Lowe and Maria Voigt and Stephen K. Burley},
   doi = {10.1002/pro.4200},
   issn = {0961-8368},
   issue = {1},
   journal = {Protein Science},
   month = {1},
   pages = {129-140},
   title = {PDB ‐101: Educational resources supporting molecular explorations through biology and medicine},
   volume = {31},
   year = {2022}
}

@misc{Voitsitskyi2024,
   abstract = {Classical protein-ligand docking has been a cornerstone technique in computational drug discovery for decades, but has reached an accuracy and performance plateau. Recently introduced Machine Learning (ML) based docking methods offer a promising paradigm shift, but their practical adoption is hampered by accuracy-to-speed trade-offs, inadequate benchmarking standards, and questionable chemical validity of predicted poses. In this study, we introduce ArtiDock - an ML-based docking technique optimized for high-throughput virtual screening applications. To evaluate ArtiDock, we developed a dedicated performance and accuracy benchmark for pocket-specific rigid protein-ligand docking, which mimics realistic industrial drug discovery scenarios and is based on the novel PLINDER dataset. We demonstrate that ArtiDock is 29-38% more accurate in comparison to leading open-source and commercial classical docking techniques such as AutoDock, Vina, and Glide, while providing a low computational cost. ArtiDock notably excels in challenging docking scenarios involving unbound protein structures and binding sites containing ions and structured water molecules. Our results show that ArtiDock could be considered as a method of choice in high-throughput virtual screening scenarios.},
   author = {Taras Voitsitskyi and Ihor Koleiev and Roman Stratiichuk and Oleksandr Kot and Roman Kyrylenko and Illia Savchenko and Vladyslav Husak and Semen Yesylevskyy and Sergii Starosyla and Alan Nafiiev},
   doi = {10.1101/2024.03.14.585019},
   month = {3},
   title = {ArtiDock: accurate Machine Learning approach to protein-ligand docking optimized for high-throughput virtual screening},
   year = {2024}
}

@misc{Boitreaud2024,
   abstract = {We introduce Chai-1, a multi-modal foundation model for molecular structure prediction that performs at the state-of-the-art across a variety of tasks relevant to drug discovery. Chai-1 can optionally be prompted with experimental restraints (e.g. derived from wet-lab data) which boosts performance by double-digit percentage points. Chai-1 can also be run in single-sequence mode with-out MSAs while preserving most of its performance. We release Chai-1 model weights and inference code as a Python package for non-commercial use and via a web interface where it can be used for free including for commercial drug discovery purposes.},
   author = {Jacques Boitreaud and Jack Dent and Matthew McPartlon and Joshua Meier and Vinicius Reis and Alex Rogozhnikov and Kevin Wu},
   doi = {10.1101/2024.10.10.615955},
   month = {10},
   title = {Chai-1: Decoding the molecular interactions of life},
   year = {2024}
}
@article{Macarron2011,
   author = {Ricardo Macarron and Martyn N. Banks and Dejan Bojanic and David J. Burns and Dragan A. Cirovic and Tina Garyantes and Darren V. S. Green and Robert P. Hertzberg and William P. Janzen and Jeff W. Paslay and Ulrich Schopfer and G. Sitta Sittampalam},
   doi = {10.1038/nrd3368},
   issn = {1474-1776},
   issue = {3},
   journal = {Nature Reviews Drug Discovery},
   month = {3},
   pages = {188-195},
   title = {Impact of high-throughput screening in biomedical research},
   volume = {10},
   year = {2011}
}
@article{Mayr2009,
   author = {Lorenz M Mayr and Dejan Bojanic},
   doi = {10.1016/j.coph.2009.08.004},
   issn = {14714892},
   issue = {5},
   journal = {Current Opinion in Pharmacology},
   month = {10},
   pages = {580-588},
   title = {Novel trends in high-throughput screening},
   volume = {9},
   year = {2009}
}
@misc{Passaro2025,
   abstract = { Accurately modeling biomolecular interactions is a central challenge in modern biology. While recent advances, such as AlphaFold3 and Boltz-1, have substantially improved our ability to predict biomolecular complex structures, these models still fall short in predicting binding affinity, a critical property underlying molecular function and therapeutic efficacy. Here, we present Boltz-2, a new structural biology foundation model that exhibits strong performance for both structure and affinity prediction. Boltz-2 introduces controllability features including experimental method conditioning, distance constraints, and multi-chain template integration for structure prediction, and is, to our knowledge, the first AI model to approach the performance of free-energy perturbation (FEP) methods in estimating small molecule–protein binding affinity. Crucially, it achieves strong correlation with experimental readouts on many benchmarks, while being at least 1000 × more computationally efficient than FEP. By coupling Boltz-2 with a generative model for small molecules, we demonstrate an effective workflow to find diverse, synthesizable, high-affinity binders, as estimated by absolute FEP simulations on the TYK2 target. To foster broad adoption and further innovation at the intersection of machine learning and biology, we are releasing Boltz-2 weights, inference, and training code 1 under a permissive open license, providing a robust and extensible foundation for both academic and industrial research. },
   author = {Saro Passaro and Gabriele Corso and Jeremy Wohlwend and Mateo Reveiz and Stephan Thaler and Vignesh Ram Somnath and Noah Getz and Tally Portnoi and Julien Roy and Hannes Stark and David Kwabi-Addo and Dominique Beaini and Tommi Jaakkola and Regina Barzilay},
   doi = {10.1101/2025.06.14.659707},
   month = {6},
   title = {Boltz-2: Towards Accurate and Efficient Binding Affinity Prediction},
   year = {2025}
}
