## Supplementary figures and images for "Accelerating Drug Discovery with HyperLab: An Easy-to-Use AI-Driven Platform"

### admet.png

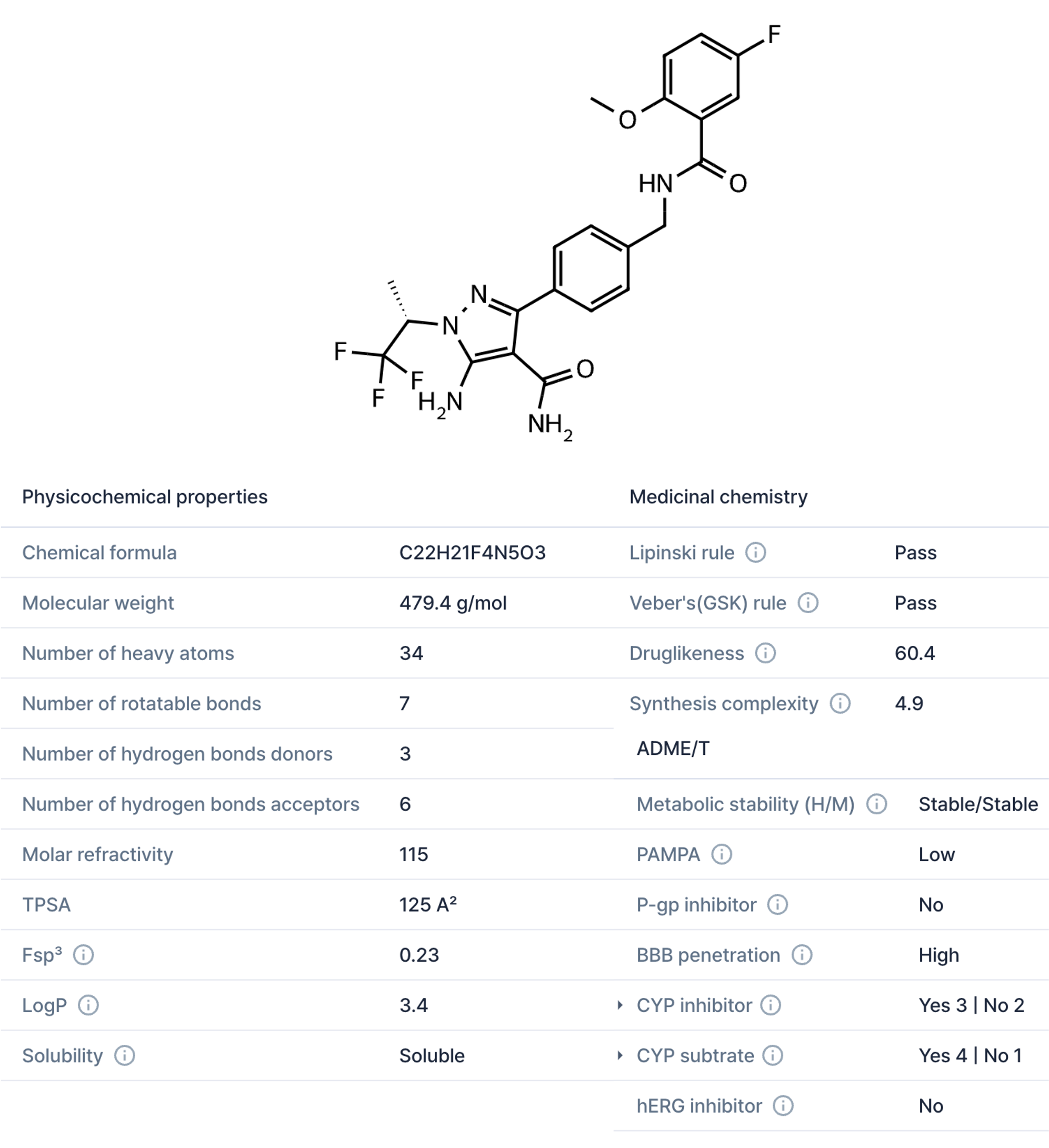

### benchmark1.png

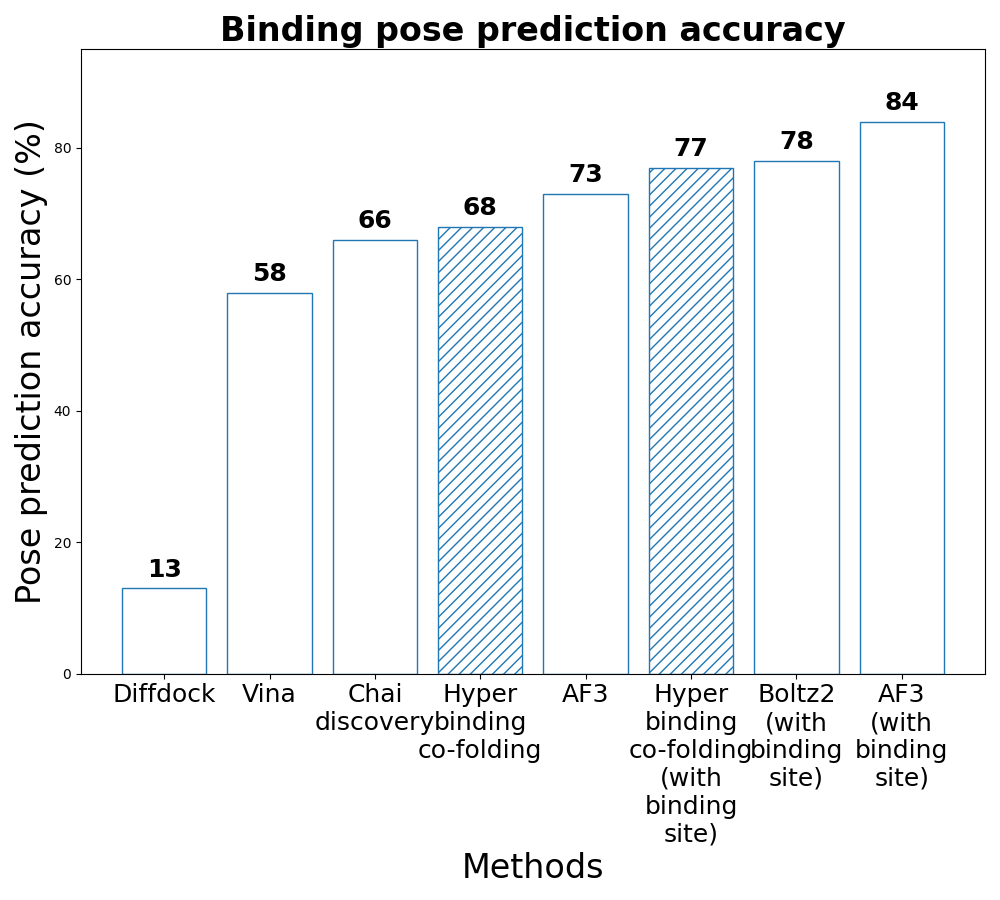

### benchmark2.png

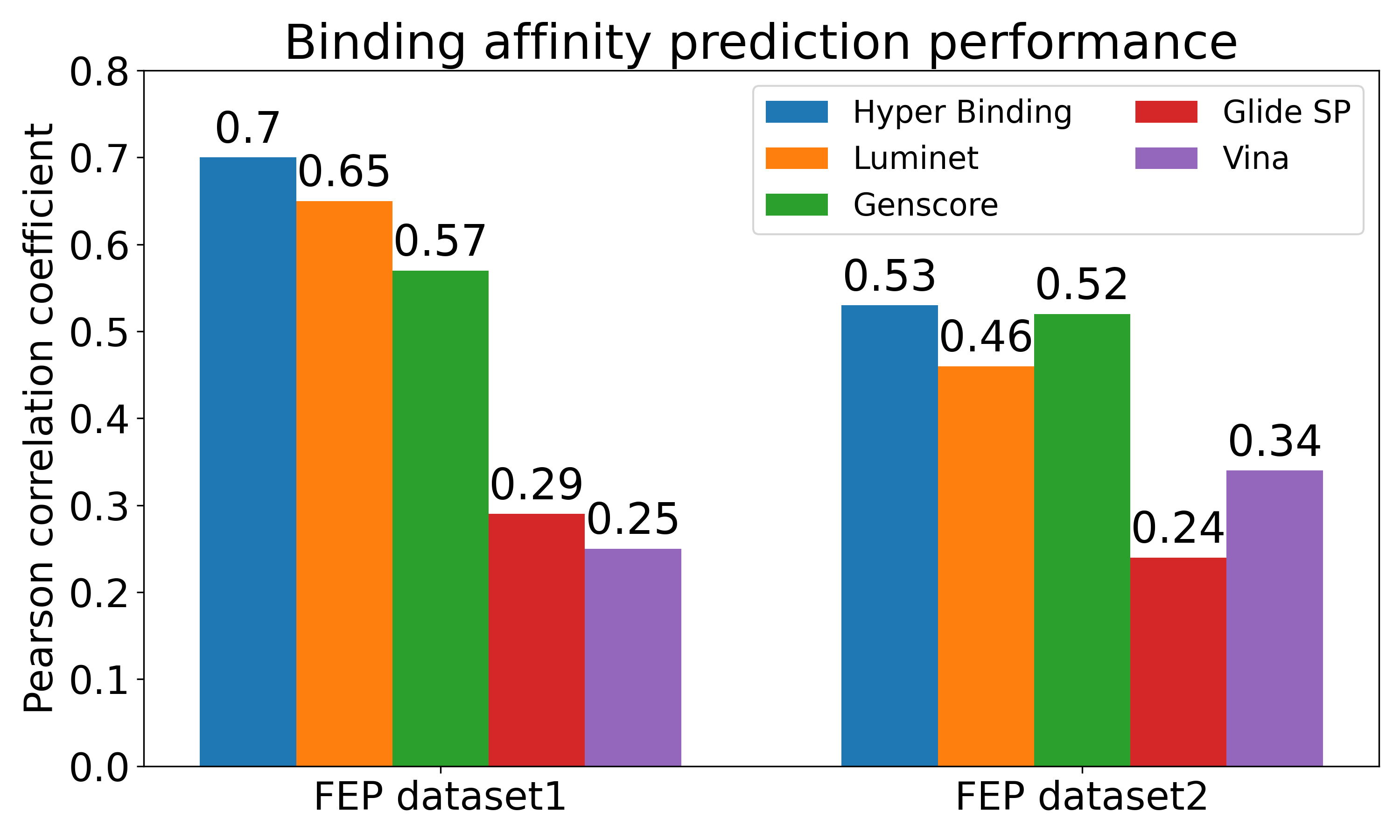

### BI2852.png

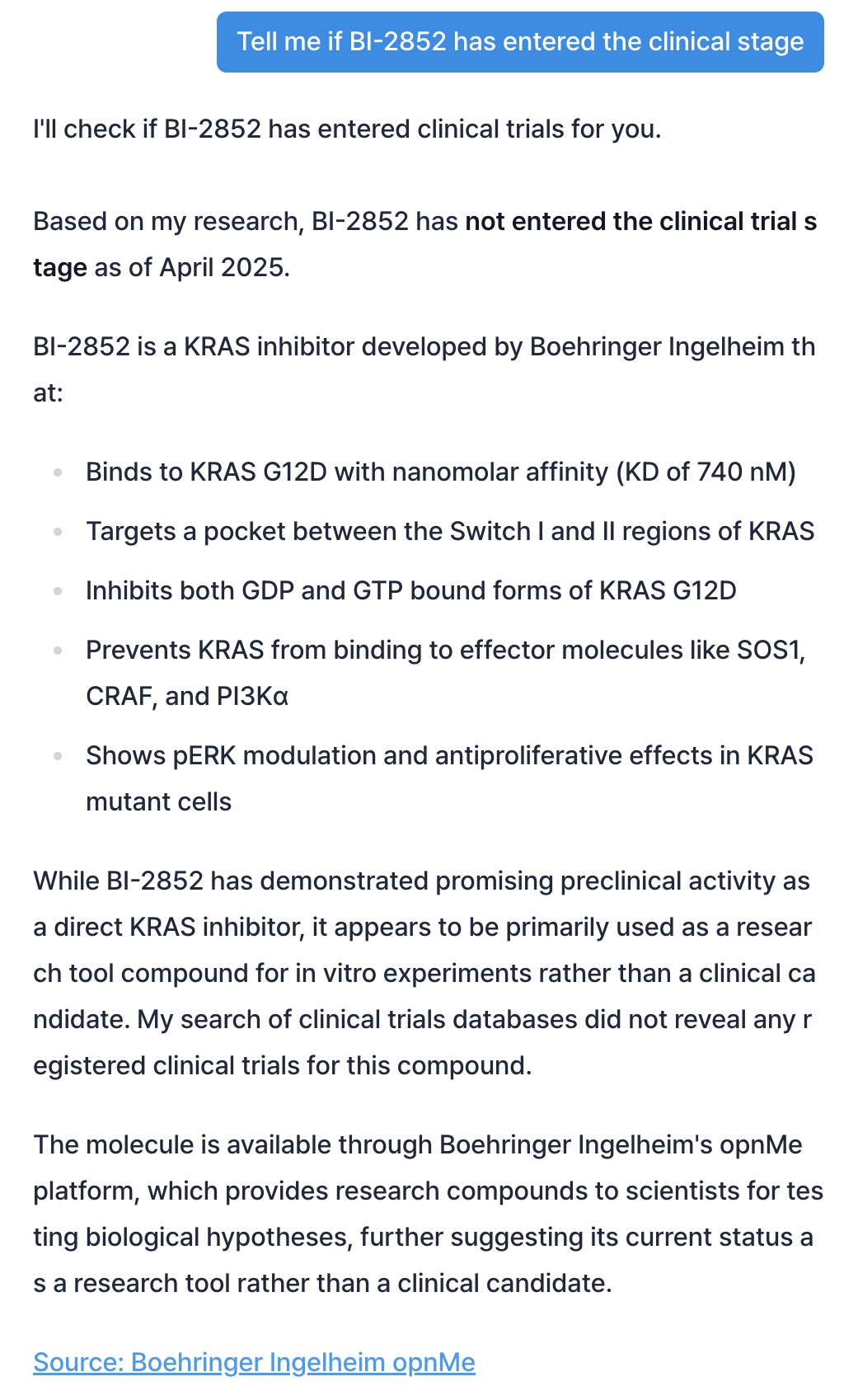

### casestudy_1.jpg

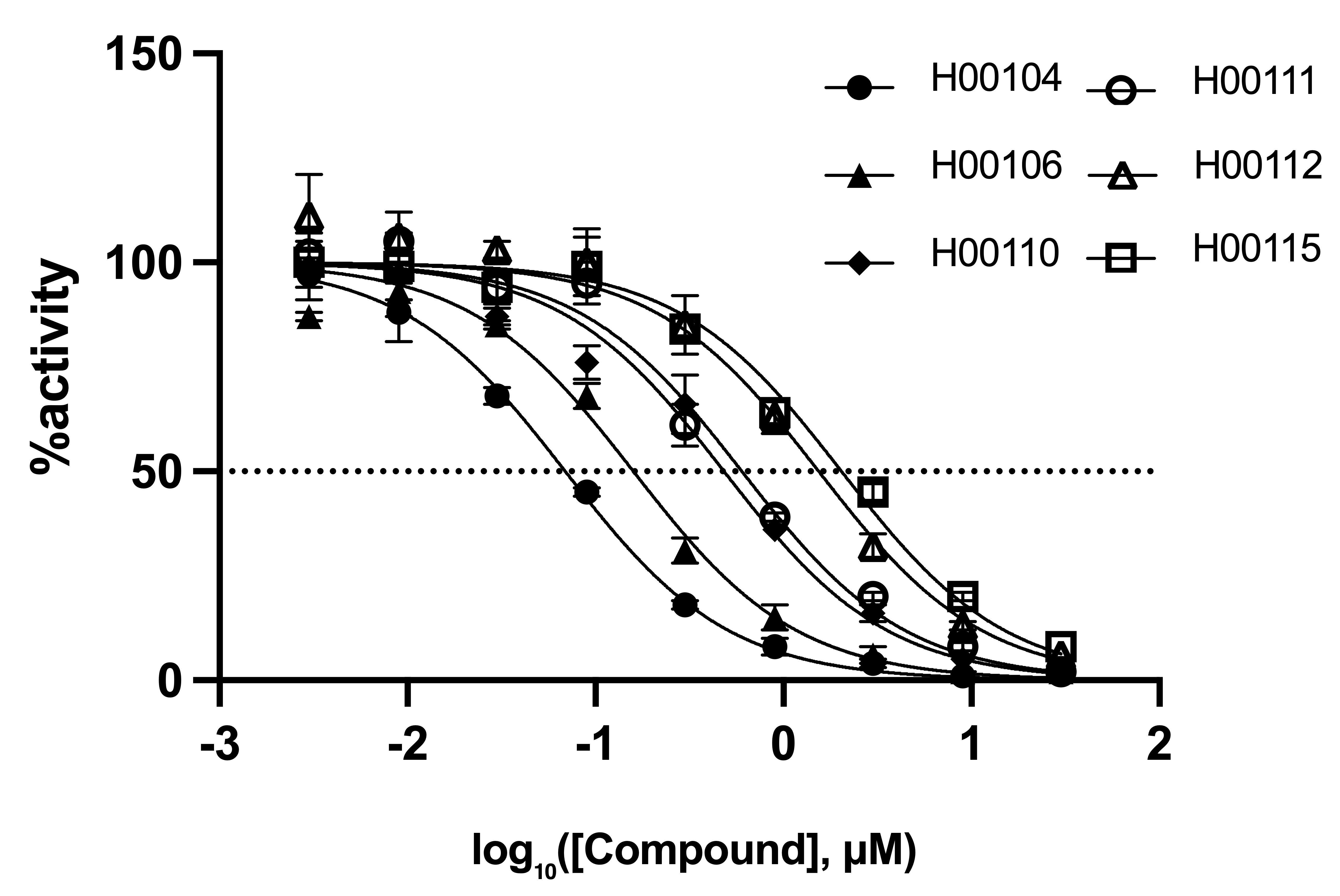

### casestudy_2.png

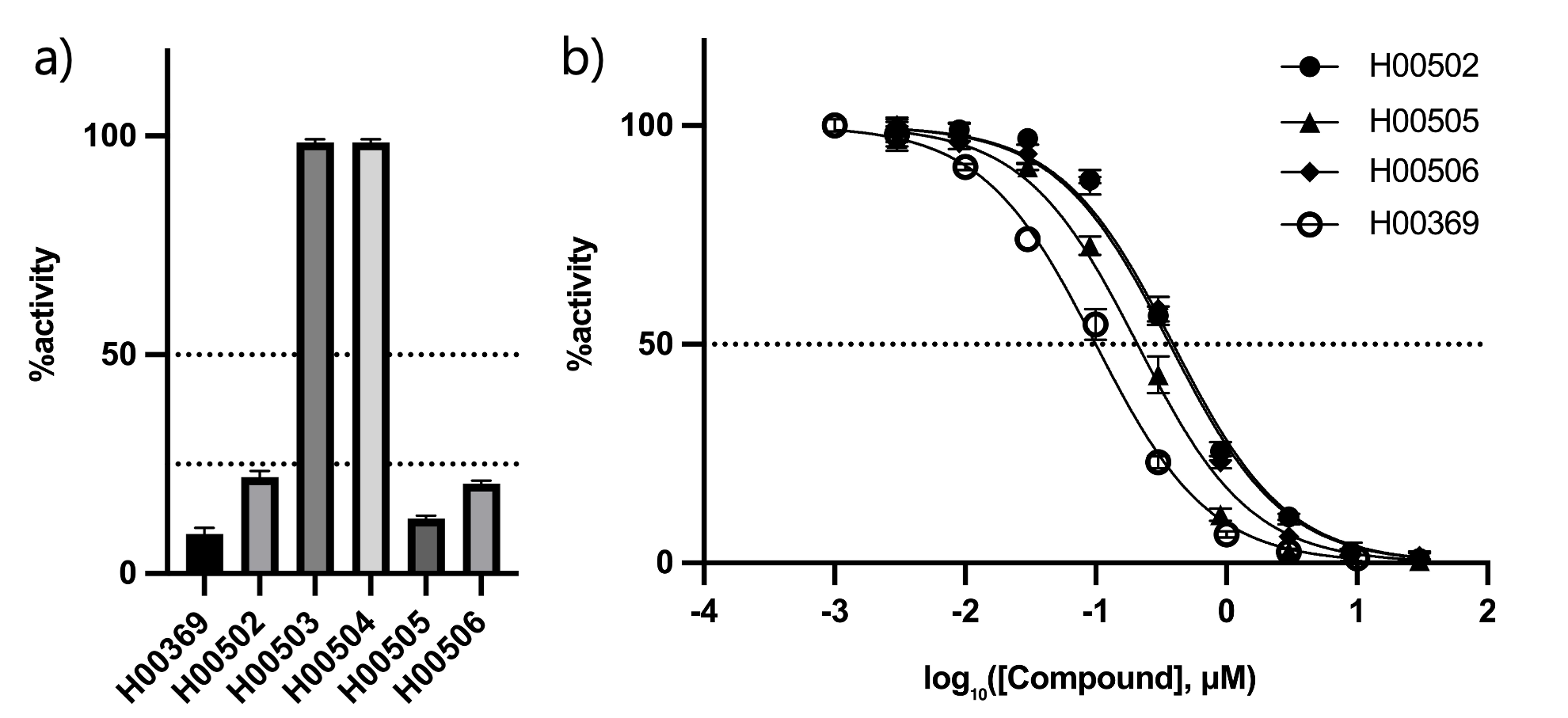

### casestudy_3.jpg

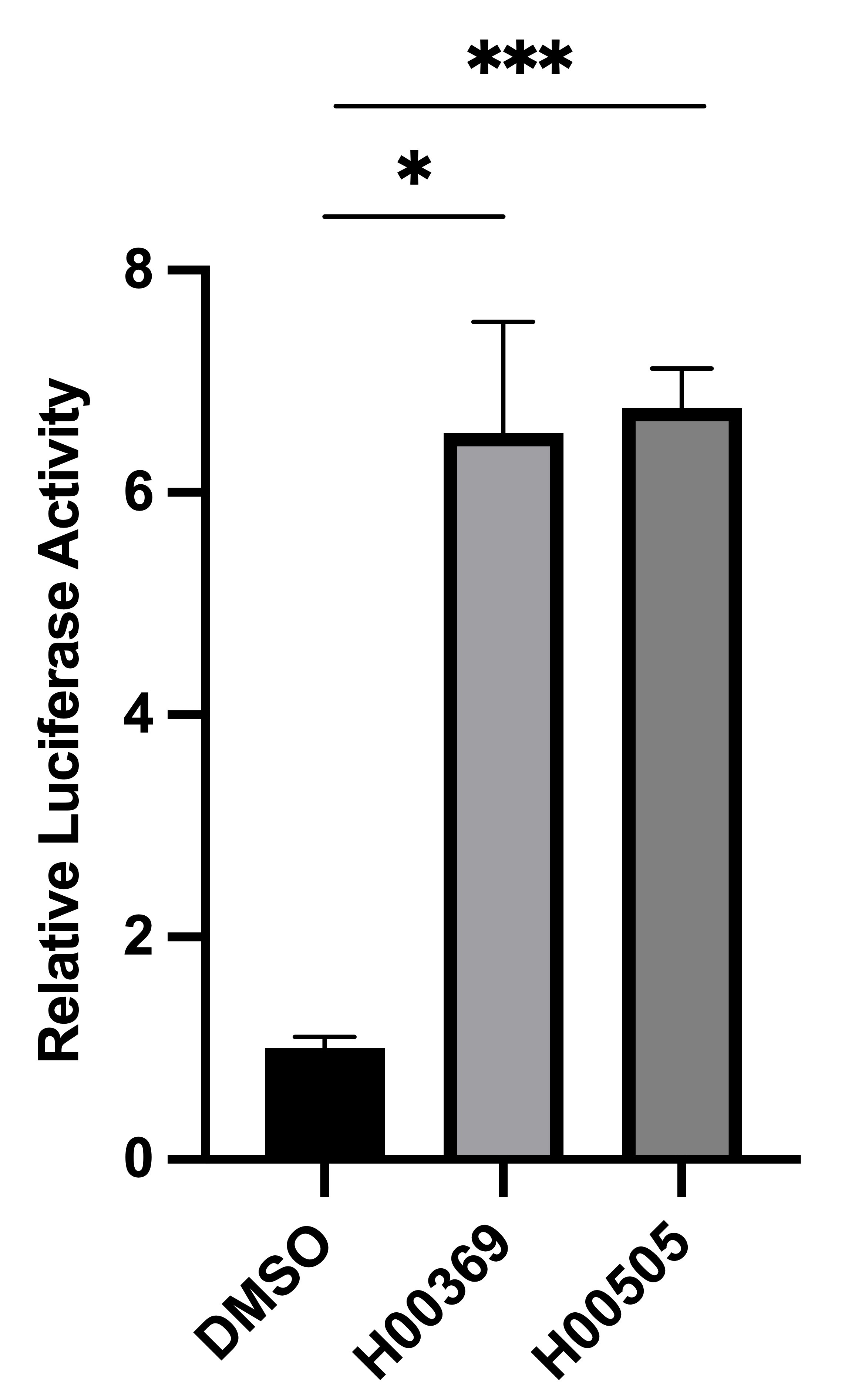

### Figure_1.png

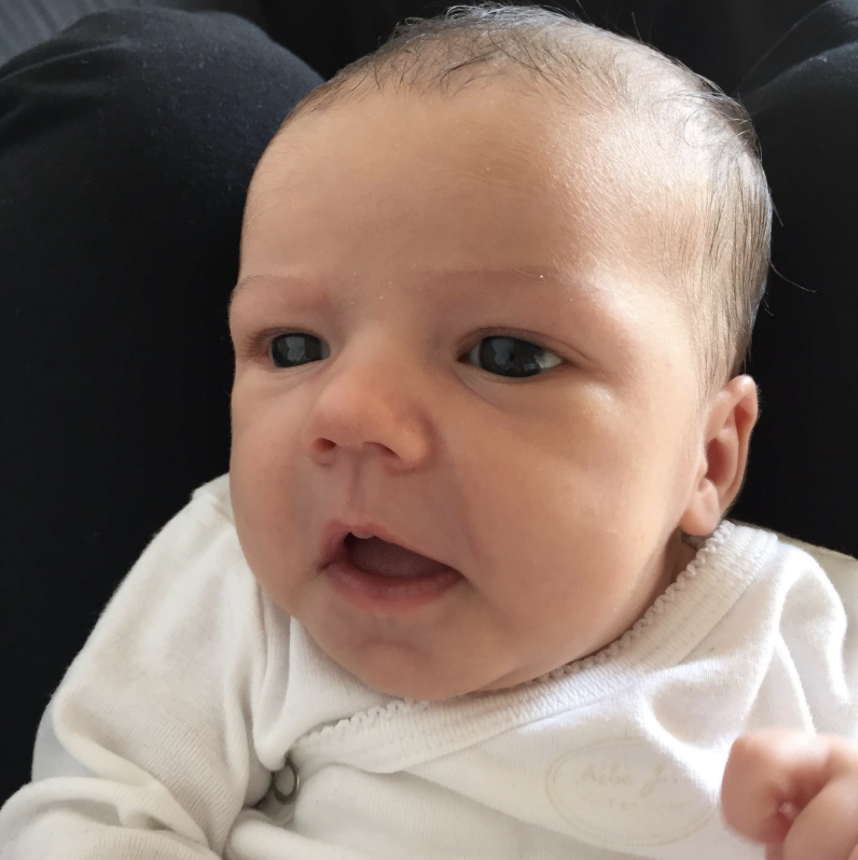

### hyperbinding.png

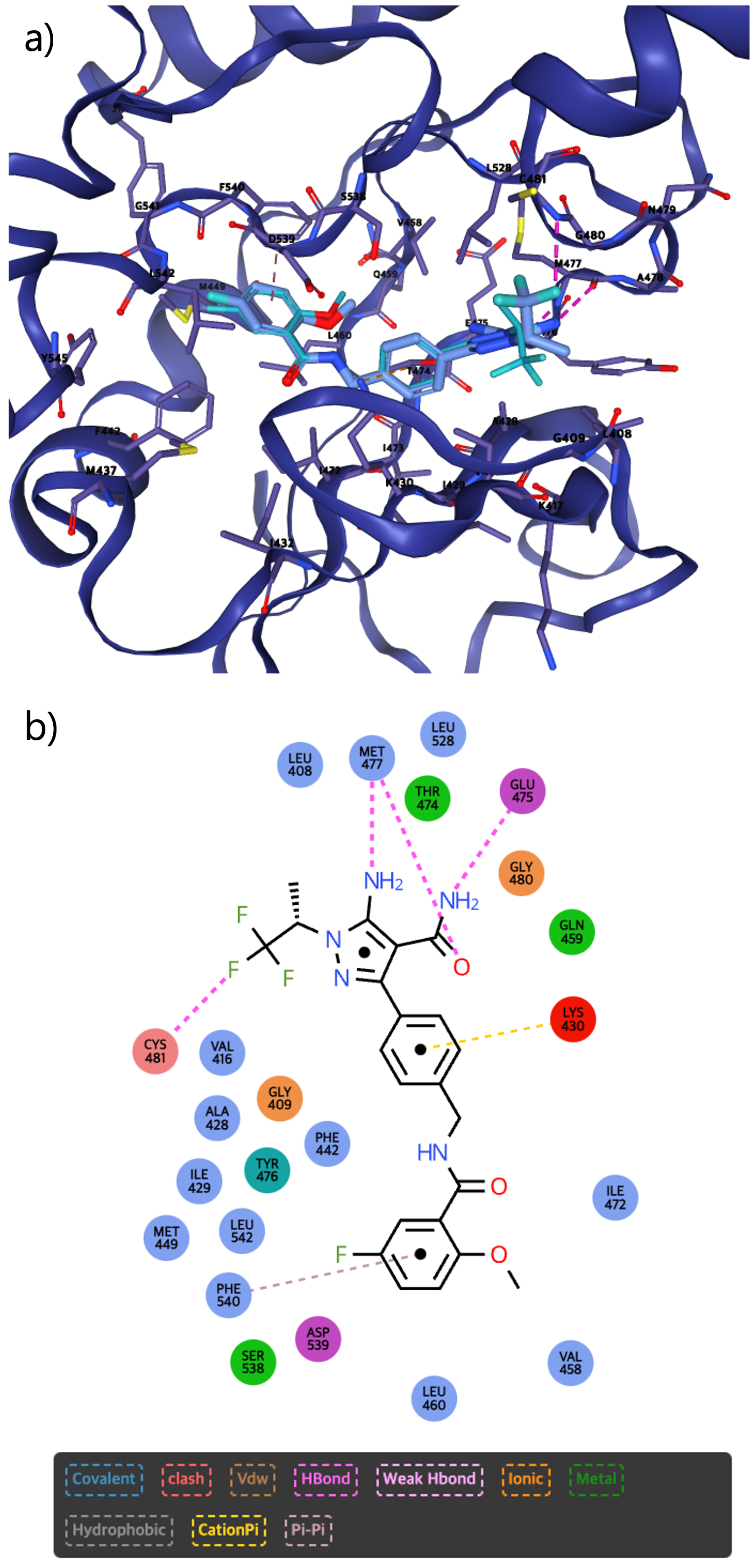

### hyperdesign.png

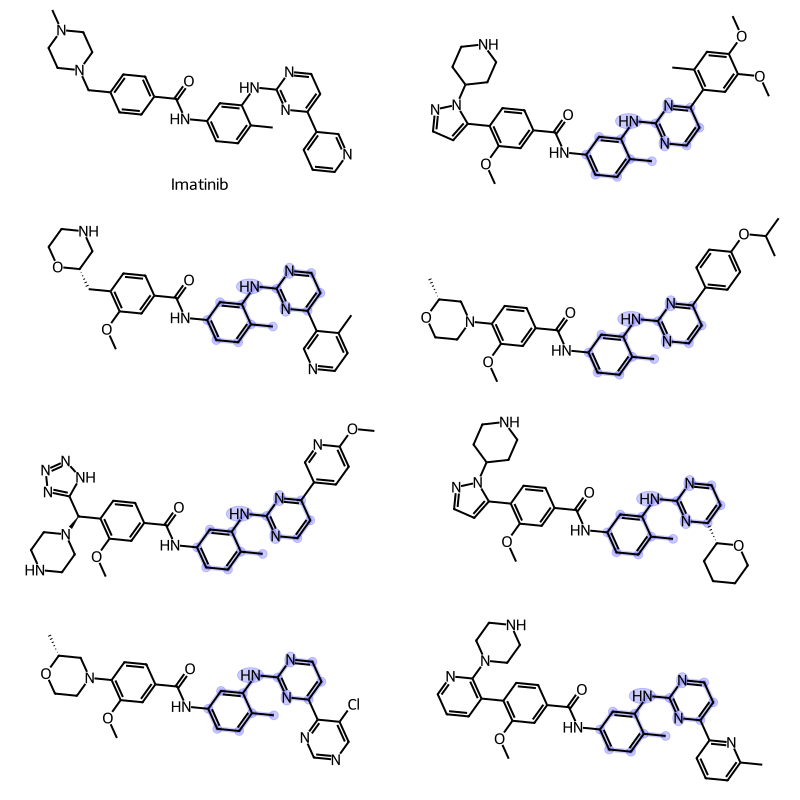

### hyperscreeningx.png

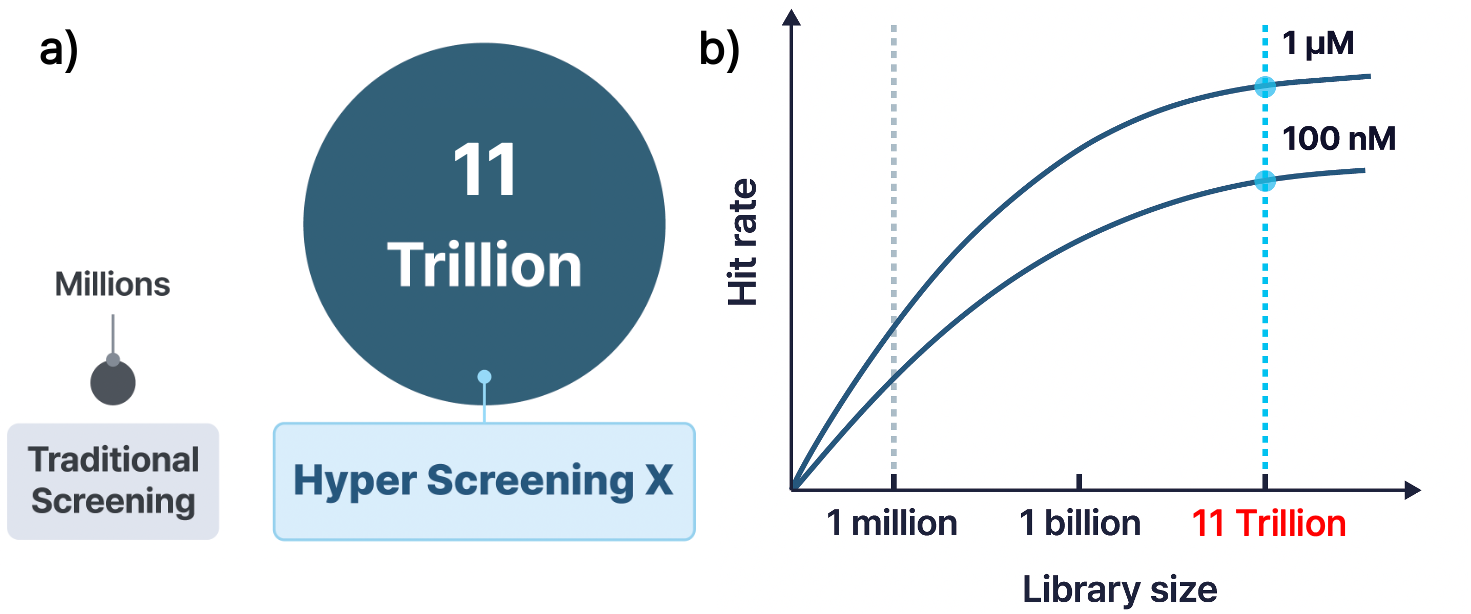

### overview.png

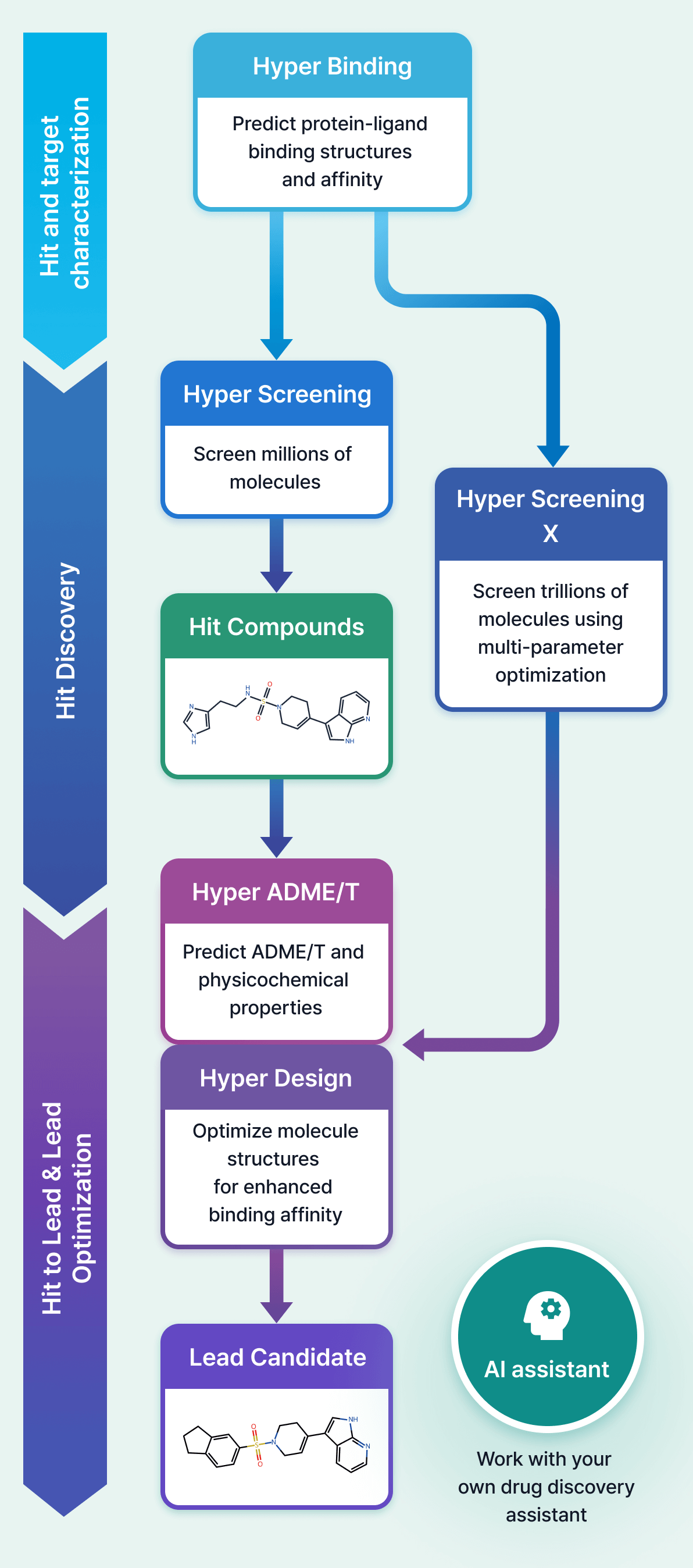

### sar.png

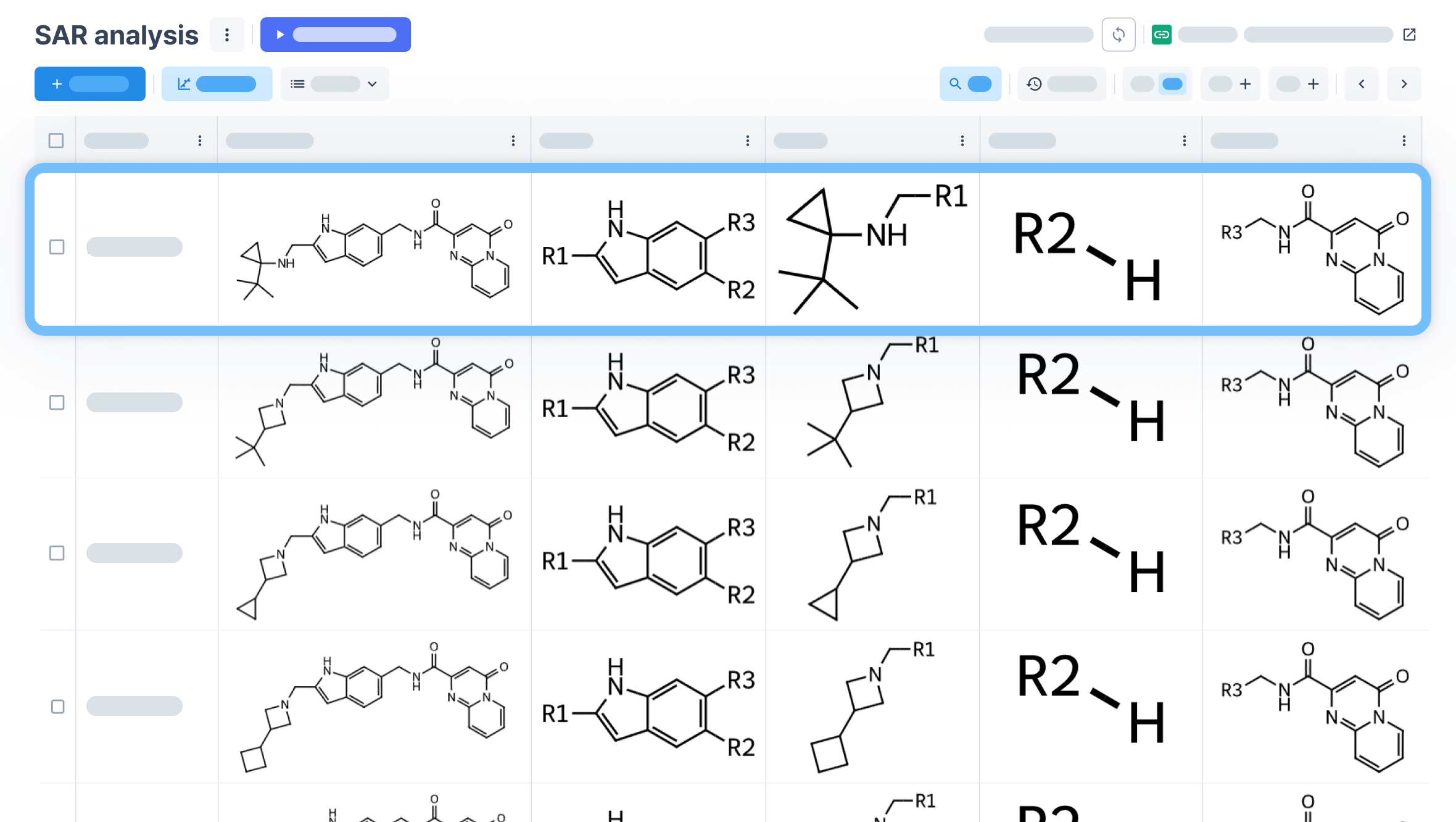
